# The dark side of the mean: brain structural heterogeneity in schizophrenia and its polygenic risk

**DOI:** 10.1101/407890

**Authors:** Dag Alnæs, Tobias Kaufmann, Dennis van der Meer, Aldo Córdova-Palomera, Jaroslav Rokicki, Torgeir Moberget, Francesco Bettella, Ingrid Agartz, Deanna M. Barch, Alessandro Bertolino, Christine L. Brandt, Simon Cervenka, Srdjan Djurovic, Nhat Trung Doan, Sarah Eisenacher, Helena Fatouros-Bergman, Lena Flyckt, Annabella Di Giorgio, Beathe Haatveit, Erik G. Jönsson, KaSP Consortium, Peter Kirsch, Martina J. Lund, Andreas Meyer-Lindenberg, Giulio Pergola, Emanuel Schwarz, Olav B. Smeland, Tiziana Quarto, Mathias Zink, Ole A. Andreassen, Lars T. Westlye

**Affiliations:** Norwegian Centre for Mental Disorders Research (NORMENT), KG Jebsen Centre for Psychosis Research, Division of Mental Health and Addiction, Oslo University Hospital, and Institute of Clinical Medicine, University of Oslo, Norway; Centre for Psychiatry Research, Department of Clinical Neuroscience, Karolinska Institutet, Stockholm, Sweden; Department of Psychological and Brain Sciences, Washington University in St. Louis, USA; Psychiatric Neuroscience Group, Department of Basic Medical Sciences, Neuroscience and Sense Organs, University of Bari “Aldo Moro”, Bari, Italy; Department of Medical Genetics, Oslo University Hospital, Oslo, Norway; Central Institute of Mental Health, University of Heidelberg, Mannheim, Germany; Fondazione Casa Sollievo della Sofferenza IRCCS, San Giovanni Rotondo, Italy; Karolinska Schizophrenia Project (KaSP) Consortium; Medical Faculty Mannheim, University of Heidelberg, Mannheim, Germany; Department of Psychology, University of Oslo, Norway; Centre for Psychiatry Research, Department of Clinical Neuroscience, Karolinska Institutet, & Stockholm County Council, Stockholm, Sweden; Department of Physiology and Pharmacology, Karolinska Institutet, Stockholm, Sweden; Neuroimmunology Unit, Department of Clinical Neuroscience, Karolinska Institutet, Stockholm, Sweden; NORMENT, KG Jebsen Centre for Psychosis Research, Division of Mental Health and Addiction, University of Oslo, Oslo, Norway; Department of Psychiatry Research, Diakonhjemmet Hospital, Oslo, Norway

## Abstract

**Importance:** Between-subject variability in brain structure is determined by gene-environment interactions, possibly reflecting differential sensitivity to environmental and genetic perturbations. Magnetic resonance imaging (MRI) studies have revealed thinner cortices and smaller subcortical volumes in patients. However, such group-level comparisons may mask considerable within-group heterogeneity, which has largely remained unnoticed in the literature

**Objective:** To compare brain structural variability between individuals with SZ and healthy controls (HC) and to test if respective variability reflects the polygenic risk for SZ (PRS) in HC.

**Design, Setting, and Participants:** We compared MRI derived cortical thickness and subcortical volumes between 2,010 healthy controls and 1,151 patients with SZ across 16 cohorts. Secondly, we tested for associations between PRS and MRI features in 12,490 participants from UK Biobank.

**Main Outcomes and Measures:** We modeled mean and dispersion effects of SZ and PRS using double generalized linear models. We performed vertex-wise analyses for thickness, and region-of-interest analysis for cortical, subcortical and hippocampal subfield volumes. Follow-up analyses included within-sample analysis, controlling for intracranial volume and population covariates, test of robustness of PRS threshold, and outlier removal.

**Results:** Compared to controls, patients with SZ showed higher heterogeneity in cortical thickness, cortical and ventricle volumes, and hippocampal subfields. Higher PRS was associated with thinner frontal and temporal cortices, as well as smaller left CA2/3, but was not significantly associated with dispersion.

**Conclusion and relevance:** SZ is associated with substantial brain structural heterogeneity beyond the mean differences. These findings possibly reflect higher differential sensitivity to environmental and genetic perturbations in patients, supporting the heterogeneous nature of SZ. Higher PRS for SZ was associated with thinner fronto-temporal cortices and smaller subcortical volumes, but there were no significant associations with the heterogeneity in these measures, i.e. the variability among individuals with high PRS were comparable to the variability among individuals with low PRS. This suggests that brain variability in SZ results from interactions between environmental and genetic factors that are not captured by the PGR. Factors contributing to heterogeneity in fronto-temporal cortices and hippocampus are thus key to further our understanding of how genetic and environmental factors shape brain biology in SZ.

**Key Points:** **Question:** Is schizophrenia and its polygenic risk associated with brain structural
heterogeneity in addition to mean changes?

**Findings:** In a sample of 1151 patients and 2010 controls, schizophrenia was associated with increased heterogeneity in fronto-temporal thickness, cortical, ventricle, and hippocampal volumes, besides robust reductions in mean estimates. In an independent sample of 12,490 controls, polygenic risk for schizophrenia was associated with thinner fronto-temporal cortices and smaller CA2/3 of the left hippocampus, but not with heterogeneity.

**Meaning:** Schizophrenia is associated with increased inter-individual differences in brainstructure, possibly reflecting clinical heterogeneity, gene-environment interactions, or secondary disease factors.

## Introduction

Schizophrenia (SZ) is a severe psychiatric disorder with a lifetime prevalence of about 1%, rendering it a leading cause of disability worldwide with 26 million people affected^1^. While genetic and environmental factors contributing to disease risk have been identified, the pathophysiology still remains elusive^2,3^. Patients diagnosed with SZ display substantial heterogeneity in terms of their clinical characteristics and symptoms^4^, treatment response^5^ and long term prognosis^6^. The notion that the observed heterogeneity stems at least partially from distinct subtypes of patients with differentially affected neurobiology and clinical and cognitive profiles^7-9^, has not been fully confirmed^10^, and the question of whether there is one unifying pathophysiological process shared across patients, or a multitude of disease processes leading to a similar clinical syndrome remains salient^11^.

SZ is associated with widespread brain abnormalities, with the most robust group-level mean structural differences being ventricle enlargement, reduced thickness and area of frontal and temporal cortices, as well as reduced hippocampal and amygdala volumes^12-14^. However there is also substantial variability between patients^7,8,15,16^, presenting a major challenge for achieving imaging based diagnostic predictions with any clinical utility^17,18^. Rather than simply reflecting noise, this inter-individual variability in brain structure may possibly carry relevant information regarding gene-environment interactions related to the individual sensitivity to environmental and genetic perturbation. Only a few studies have investigated whether heterogeneity differs between healthy participants and SZ patients. One functional imaging study reported increased heterogeneity in both connectivity and spatial extent of functional brain networks in SZ^19^. Regions with altered spatial variance in functional networks included areas previously implicated in SZ, such as auditory and sensorimotor cortices and basal ganglia, and networks showing increased heterogeneity overlapped with those showing mean volume differences, implying that the mean and variance measures provide complementary but converging results^20^. A recent meta-analysis reported increased inter-individual volumetric variability in several cortical and subcortical structures, including the temporal lobe, thalamus, hippocampus and amygdala in SZ, and lower variability in the anterior cingulate cortex (ACC)^15^. These results point to the importance of modeling heterogeneity as well as mean changes. Detecting brain regions that are more homogenous in patients could point to a primary role in a shared underlying SZ pathophysiology, while regions of increased heterogeneity might be informative of putative subtypes of disease, or possibly reflect regional differences in the sensitivity to genetic and environmental perturbations.

SZ is highly heritable^21^, motivating the ongoing efforts to identify intermediate brain phenotypes associated with disease liability in order to elucidate the pathway from genes to illness manifestation. Several SZ risk loci have been identified^22^, but the individual contribution of each identified variant is weak and as of yet no common variants have been conclusively linked to the disease. Polygenic risk scores (PRS) for SZ, which represent a weighted sum of common genetic SZ risk alleles, have been proposed to account for the polygenic nature of disease risk and to increase predictive power^23^. Beyond being predictive of case-control status^22^, SZ PRS have been associated with negative symptoms, anxiety and lower cognitive ability in adolescents^24^. Polygenic burden has also been linked to reduced cortical thickness, as well as with prefrontal working memory-, and hippocampal encoding-related activation and connectivity in both patients and healthy participants^25-28^. This is in line with findings implicating both the frontal cortices and hippocampus as core regions in SZ pathophysiology^29^. Polygenic risk for SZ is however only weakly associated with subcortical volumes^30^. Importantly, risk alleles could also exert their effect by influencing the environmental sensitivity, which could be reflected in the phenotypic variability between individuals^31^.

Thus, revealing brain structures with higher or lower heterogeneity in SZ could facilitate discovery of intermediate brain phenotypes that may serve to identify putative sub-types^8,32^ of the disease, as well as by identifying phenotypes that are primary or common in the neurobiology of SZ^15^. Further, investigating how the genetic architecture of disease risk is related to brain heterogeneity could reveal regions in which the cumulative burden of common risk alleles influence the phenotypic variance^33^. To this end, we directly compared the within-group dispersion in several key brain structural phenotypes, including cortical thickness, as well as in cortical, subcortical and hippocampal subfield volumes, between 1151 patients with SZ and 2010 healthy controls. Next, in order to test the whether between-subject variability is associated with the cumulative polygenic risk for SZ, we tested for associations between dispersion in the same brain features and PGR for SZ in 12,490 healthy individuals from the UK Biobank.

## Material and methods

### Samples

Sample characteristics are presented in Table 1 and eMethods (Samples), and cohort details are presented in eTable 1.

**Table 1.** Sample characteristics. We included data from 16 samples for the case-control analysis, each sample contained MR-scans from both patients with SZ and healthy control participants. -analysis was performed on UKB participants, excluding those with a diagnosed neurological or mental disorder. More detailed information about samples and acknowledgements are provided in eTable 1.

| Case-control samples |  |  |  |  |  |  |  |  |
| --- | --- | --- | --- | --- | --- | --- | --- | --- |
| Sample | HC |  |  |  | SZ |  |  |  |
|  | n | mean age | std | females (n) | n | mean age | std | females (n) |
| BrainGluSchi | 47 | 33,7 | 12,7 | 18 | 37 | 37,2 | 15,3 | 3 |
| CIMH | 43 | 30,6 | 11,5 | 14 | 53 | 31,2 | 8,6 | 17 |
| CNP-1 | 102 | 31,7 | 8,9 | 47 | 25 | 36,9 | 9,3 | 8 |
| CPN-1 | 23 | 30,7 | 8,6 | 12 | 25 | 36,0 | 8,6 | 4 |
| COBRE | 90 | 38,1 | 11,7 | 26 | 88 | 38,1 | 13,7 | 18 |
| HUBIN | 102 | 42,0 | 8,8 | 33 | 94 | 41,7 | 7,6 | 24 |
| KaSP | 45 | 25,7 | 5,2 | 21 | 37 | 28,5 | 7,5 | 12 |
| MCIC-1 | 43 | 29,0 | 11,3 | 9 | 35 | 32,6 | 12,4 | 7 |
| MCIC-2 | 26 | 33,1 | 12,4 | 11 | 32 | 32,4 | 10,3 | 8 |
| MCIC-3 | 24 | 40,8 | 9,4 | 9 | 33 | 38,3 | 10,0 | 8 |
| NUNA | 44 | 31,3 | 8,5 | 22 | 44 | 33,0 | 6,8 | 14 |
| NUSDAST | 95 | 33,8 | 14,5 | 44 | 114 | 35,0 | 13,1 | 38 |
| TOP-3T-1 | 294 | 31,8 | 7,6 | 121 | 112 | 28,0 | 8,1 | 38 |
| TOP-3T-2 | 301 | 35,8 | 10,1 | 141 | 110 | 33,0 | 9,9 | 44 |
| TOP-1.5 | 295 | 35,5 | 9,6 | 131 | 225 | 31,9 | 9,1 | 96 |
| UNIBA | 436 | 26,7 | 7,7 | 225 | 87 | 34,1 | 7,6 | 22 |
| **Total** | **2010** | **32,7** | **10,4** | **884** | **1151** | **33,8** | **10,6** | **361** |
| PRS sample |  |  |  |  |  |  |  |  |
| **Sample** | **n** | **mean age** | **std** | **females (n)** |  |  |  |  |
| UKB | 12490 | 55,9 | 7,5 | 6465 |  |  |  |  |

### Image preprocessing

T1-images where processed using Freesurfer 5.3.0 (http://surfer.nmr.mgh.harvard.edu) for cortical reconstruction and volumetric segmentation^34-37^ and Freesurfer 6.0 for hippocampus subfield segmentation^38^. Each participant’s cortical thickness map was registered to the MNI305 fsaverage template and spatially smoothed (15 mm FWHM).

### Genetic data

PRS were calculated using PRSice^39^ v1.25, based on the European-Caucasian subset of the 2014 PGC2 schizophrenia GWAS^22^. The scores were calculated at a range of p-value thresholds, from 0.001 to 0.5, with intervals of 0.001 (eMethods: Genetics). PRS based on a threshold of .05 were used for the main analysis since this threshold has been reported as optimal in terms of explaining case-control differences^40^, but we also performed follow-up analysis to test the robustness of findings to threshold selection.

### Statistical analysis

Statistical analyses were performed in R (3.2.2; https://www.r-project.org/) and MATLAB (2014a, The MathWorks, Inc., Natick, Massachusetts, United States). For all included measures in the multi-scanner case-control sample we used vertex/volume-wise generalized additive models (GAM, R-Package: mgcv^41^) to regress out scanner effects while accounting for age, sex and diagnosis. We then modeled vertex/volume-wise mean and dispersion effects using double generalized linear models (DGLMs), which iteratively fits a GLM modeling the mean parameter, and a second GLM modeling the dispersion parameter (eMethods: DGLMs; R-package: dglm^42^). We then permuted the SZ and PRS column vector, respectively, and recalculated the mean and dispersion effects. For cortical thickness the true test statistics and the permuted statistical maps where then submitted to PALM^43^ to correct for multiple comparisons using threshold-free cluster enhancement (TFCE) and tail approximation^47^ (600 permutations: eMethods: Permutation). Maps were thresholded at a significance level of p<.05. For the cortical, subcortical, and hippocampal subfields volumes we performed 5000 permutations per volume and extracted the maximum t-value across ROIs when computing p-values in order to correct for multiple comparisons. Significance threshold was set at p<.05. We also performed a meta-analysis of the multi-scanner thickness data (R-Package: metafor^44^) estimating case-control difference within each sample, and conducted analyses both with and without covarying for estimated intracranial volume (eTIV) for the volumetric measures. We performed follow-up analyses with more stringent exclusion criteria (eMethods: Outliers). To assess the effect of PRS p-threshold selection, we performed a principal component analysis (R-package: prcomp, factoextra^45^) on PRS scores calculated across several thresholds (eMethods: PRS-PCA), and reran cortical thickness analysis on PCA-scores.

## Results

### Vertex-wise thickness

SZ was associated with decreased mean thickness globally, with the exception of the visual cortex, as well as globally increased thickness dispersion (Figure 1, panel A and B, Figure 2A; eFigure 2). Meta-analysis of within-sample effects with more stringent exclusion criteria revealed significantly increased heterogeneity in SZ, indicating that dispersion effects are not simply explained by multi-site variability or a few extreme values (eFigure 3). PRS was associated with lower mean thickness in the right inferior frontal gyrus, the right lateral orbitofrontal cortex, the right pre-central gyrus, the right medial temporal cortex, and bilaterally in middle and superior temporal cortices (Figure 1, panels C and D; eFigure 4). Converging results were obtained upon re-analysis with the addition of the first four population components added as covariates (eFigure 5A), or with more stringent exclusion criteria (eFigure 5B). Follow-up analysis using the first component from the PRS principal component analysis (PRS-PC1) gave close to an identical pattern as the PRS-model based on a threshold of .05 (eFigure 5C; vertex-wise r=.91). There was no significant associations between polygenic risk and thickness dispersion, or between PRS-PC2 or PRS-PC3 and mean or dispersion of cortical thickness. Vertex-wise correlations between raw t-maps for SZ and polygenic risk were r=.2 and r=.1, for the mean and dispersion respectively (eFigure 6).

**Figure 1.**
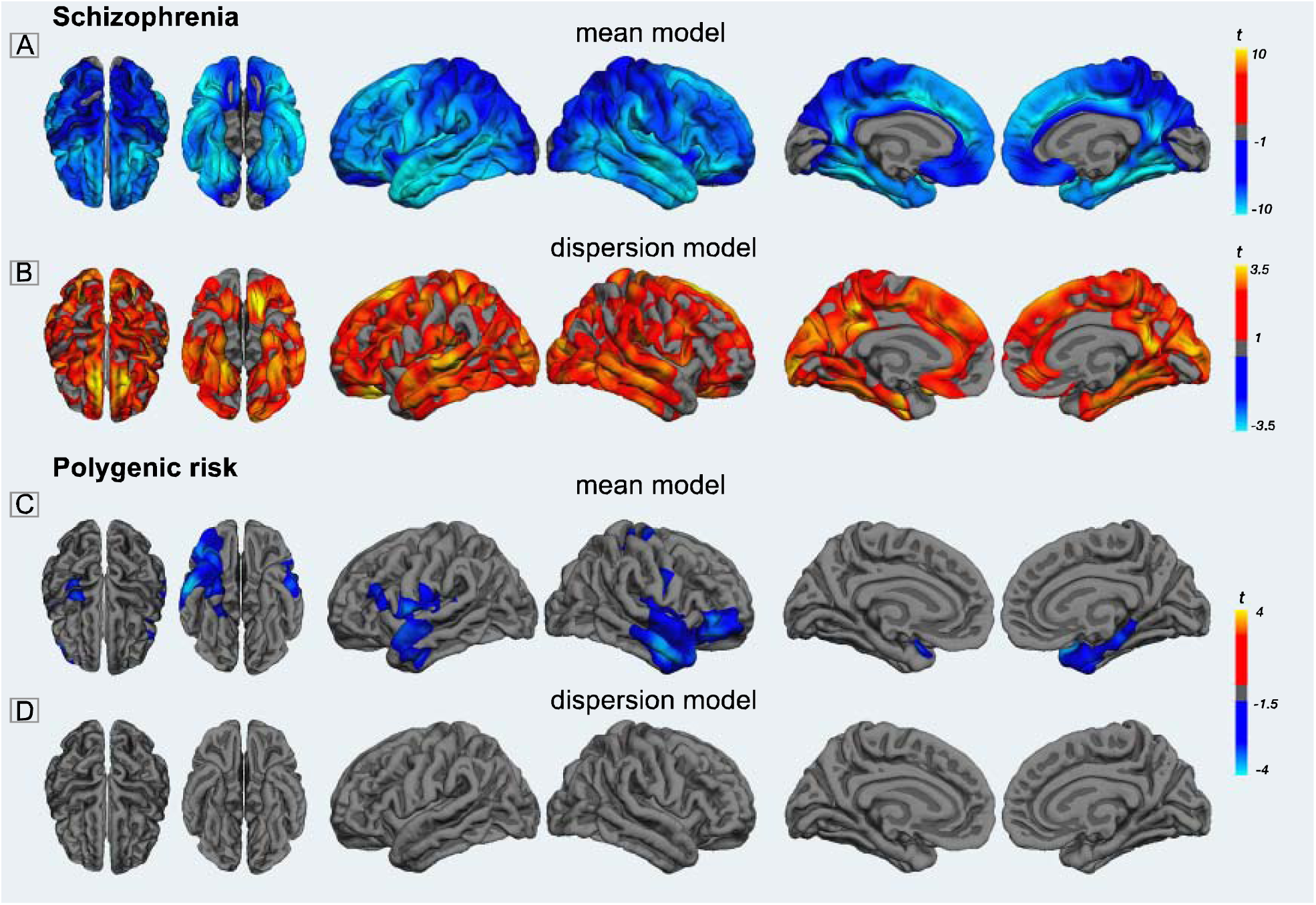
Mean and dispersion of cortical thickness. All maps were thresholded using permutation testing, threshold free cluster enhancement, and fitting the tail of the permutation distribution to a generalized Pareto distribution (500 permutations, p < .05, FWE)**. A:** t-map for the SZ mean model; cold colors represent areas with decreased mean thickness in SZ compared to healthy controls. SZ was associated with decreased thickness globally, with the exception of the visual cortex, and with strongest effects in frontal and temporal regions, compared to healthy controls. **B:** t-map for the SZ dispersion model. Warm colors represent areas with increased heterogeneity in SZ compared to healthy controls. Inter-individual variability in cortical thickness showed a spatially global increase for the SZ-group compared to healthy controls. **C:** In an independent sample of healthy adults, the mean model showed that higher polygenic risk for SZ was associated with lower cortical thickness, represented by cold colors, in frontal and parietal cortices. **D:** Polygenic risk was not associated with cortical thickness heterogeneity in any region.

**Figure 2.**
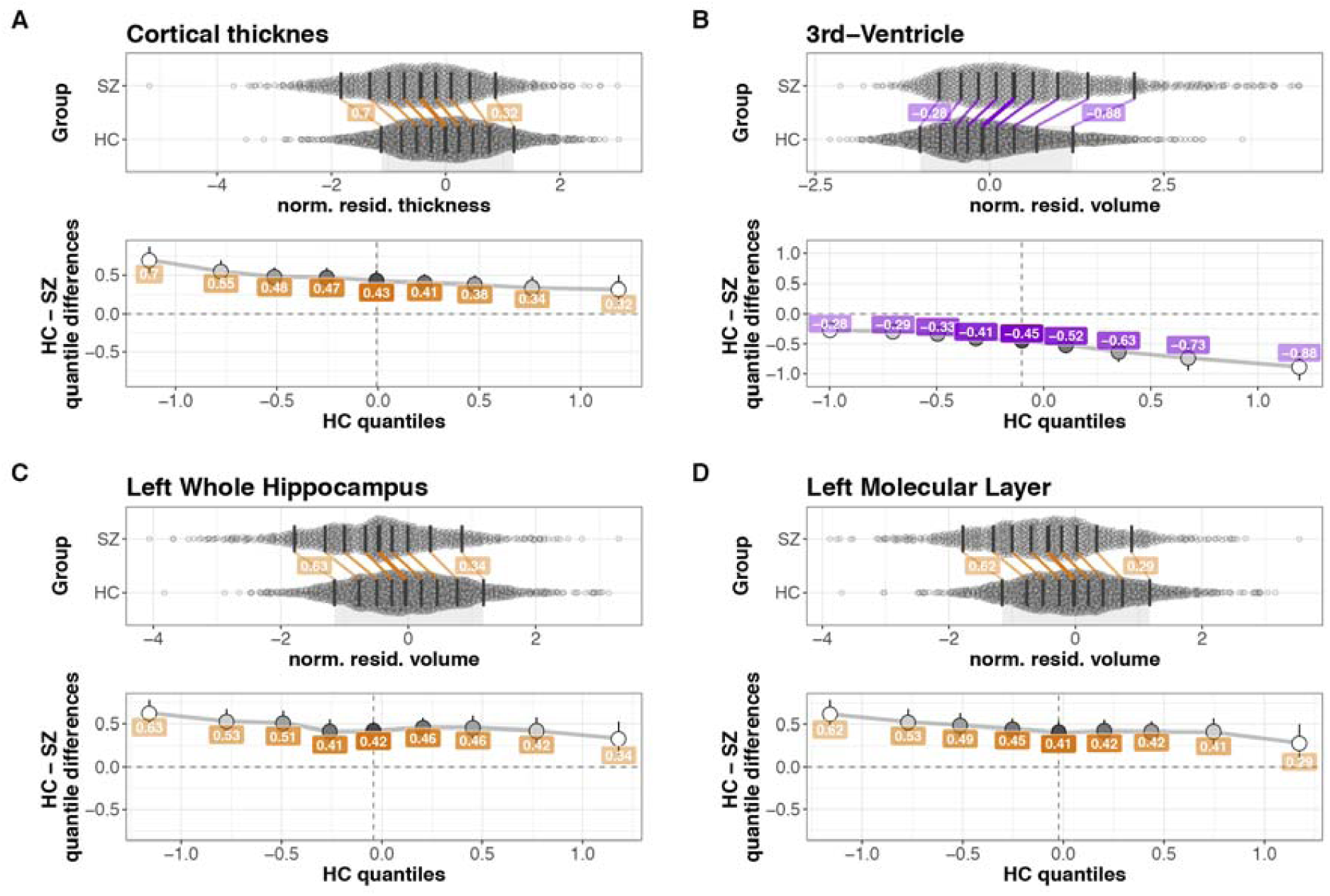
Shift function plots. The top panels show jittered marginal distribution scatterplots for variance model associations, with overlaid shift function plots using deciles. 95% confidence intervals were computed using a percentile estimation of the standard error of the difference between quantiles on 1000 bootstrap samples. The bottom panels show linked deciles from shift functions in top panels. **A**: Cortical thickness, vertex values were extracted by masking the images by the SZ-dispersion significance map, and averaged across vertices and hemispheres. Residualized for scanner, sex and age. SZ is associated with reduced thickness, with larger differences between groups in the lower deciles. **B**: 3rd ventricle volume, residualized for scanner, sex and age and eTIV. SZ is associated with larger volumes compared to controls, with the largest difference between groups in the upper deciles. **C**-**D**: Left whole hippocampus, and left molecular layer (hippocampal subfield), values residualized for scanner, sex and age and eTIV. SZ is associated with larger volumes compared to controls, with the largest difference between groups in the upper deciles.

**Figure 3.**
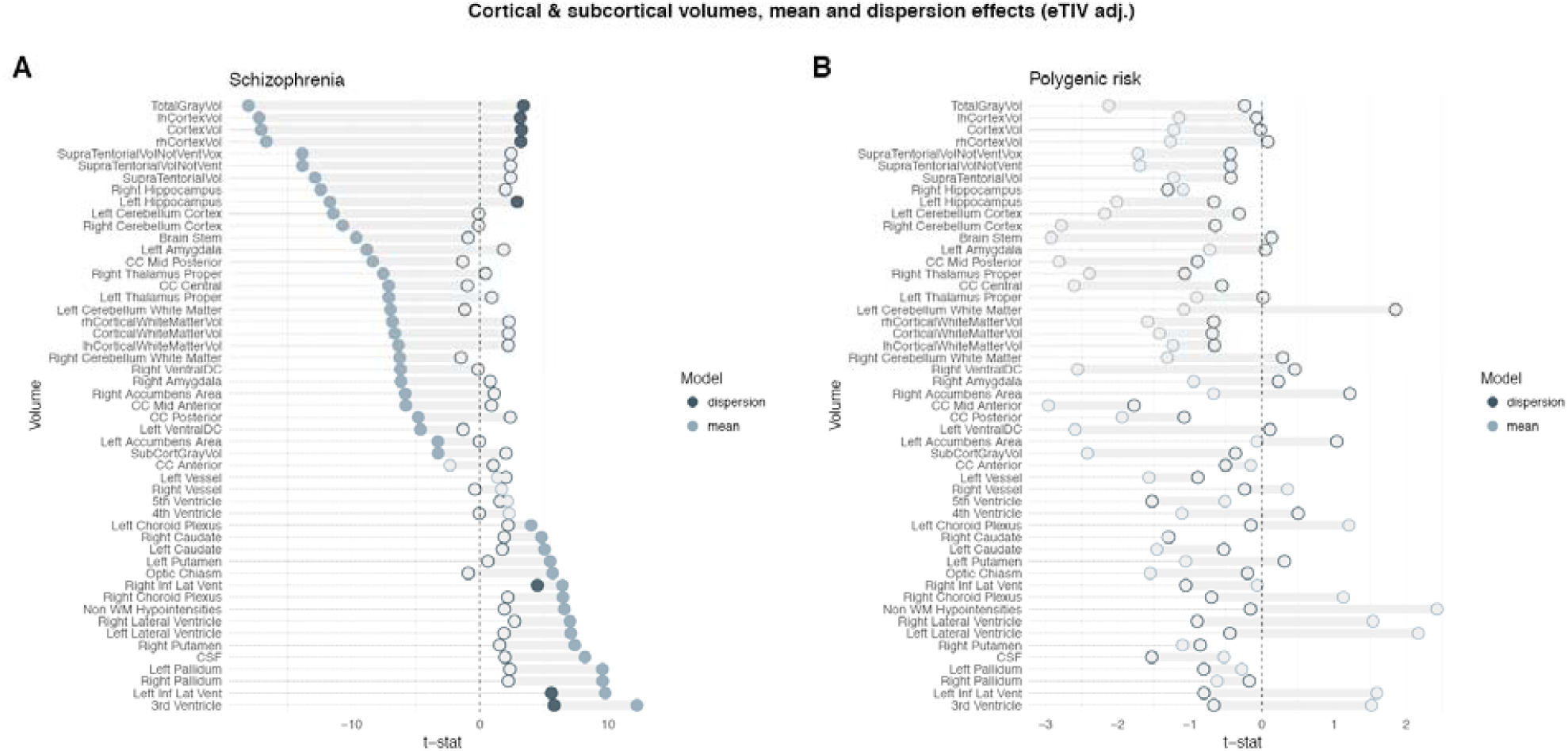
Mean and dispersion of cortical and subcortical volumes. t-statistic for both mean (outline in light blue) and dispersion (outline in dark blue), filled blue dots mark significant effects after correction for multiple comparisons across regions (5000 permutations, permuted p < .05, FWE, eTIV-adjusted). **A:** The SZ-group had both decreased cortical and subcortical volumes, as well as increased ventricles and putamen and pallidum volumes. Cortical, hippocampal and ventricle volume were more heterogeneous in the SZ-group compared to healthy controls. **B:** Polygenic risk for SZ was not associated with mean changes nor dispersion in any of the regions.

**Figure 4.**
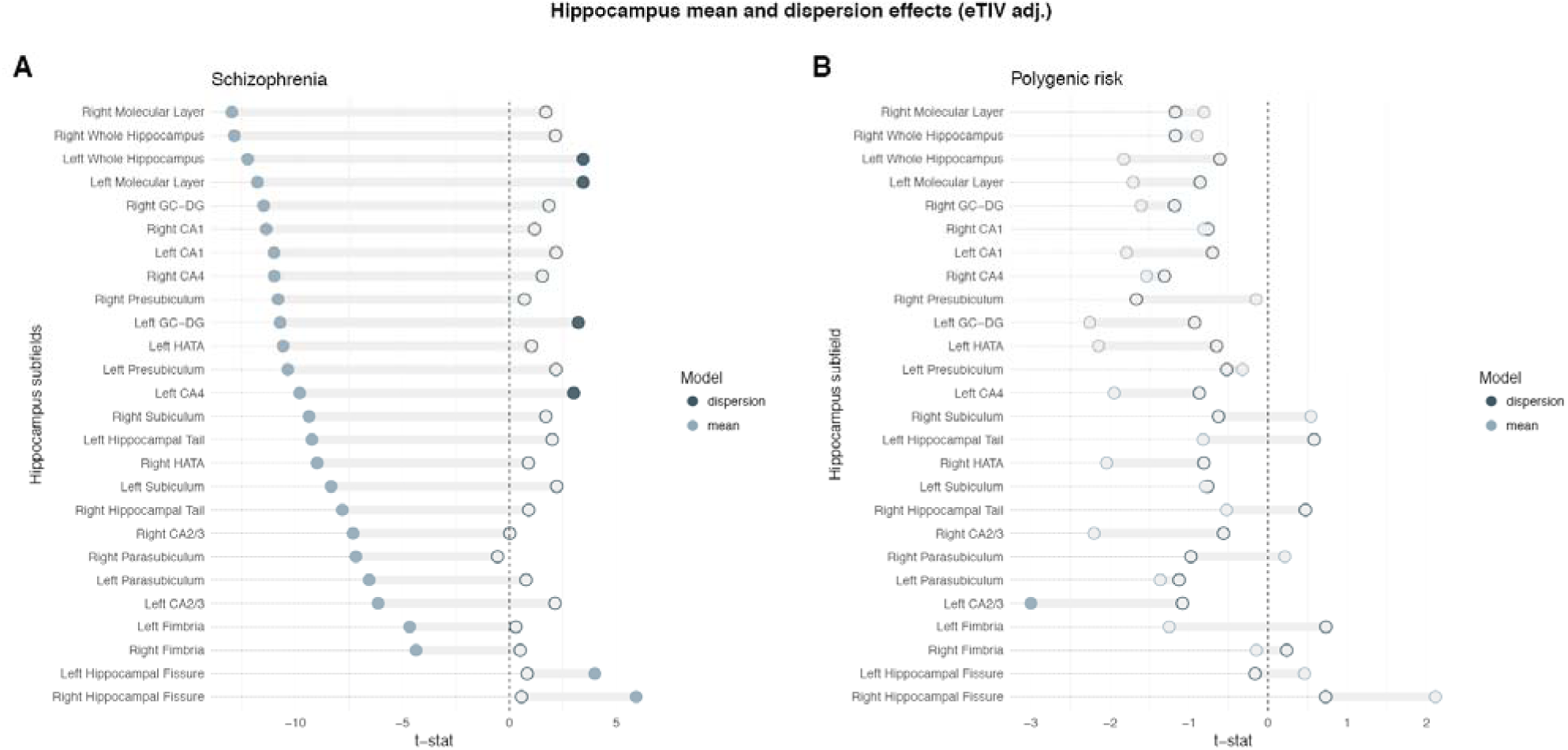
Mean and dispersion of hippocampus subfields volumes. t-statistic for both mean (outline in light blue) and dispersion (outline in dark blue), filled blue dots mark significant effects after correction for multiple comparisons across regions (5000 permutations, permuted p < .05, FWE, eTIV-adjusted). **A:** The SZ-group had decreased hippocampal volumes, and this decrease was also evident in all subfields, and accompanied with an increase of the hippocampal fissures. Hippocampal volumes were also more heterogeneous in the SZ-group, and among the subfields this effect was present in the left molecular layer, bilateral CA1, right dentate gyrus and left CA4. **B:** Polygenic risk for SZ was associated with mean reductions of left dentate gyrus, left CA4 and bilateral CA2/3. Neither total hippocampal volumes nor any of the subfields showed a significant association between polygenic risk and volume heterogeneity.

### Cortical and subcortical volumes

SZ was associated with lower mean cortical volume, supratentorial volume, total and subcortical gray volume, cerebellar cortical volume, as well as brain stem, hippocampus, amygdala, thalamus and nucleus accumbens, and several white matter volumes, as well as increased ventricle, caudate nucleus, pallidum and putamen volumes. SZ was further associated with increased dispersion in mean cortical volume, total gray volume, left hippocampus and ventricle volumes (Figure 2B, Figure 3). Models without eTIV (eFigure 7A) revealed no significant differences in the mean volumes of caudate nucleus and left putamen, and resulted in an additional significant association with dispersion in supratentorial volume. Reanalysis of mean and dispersion models with more stringent exclusion criteria showed converging results (eTable 2). PGR was not associated with mean or dispersion in any of the subcortical volumes (Figure 3B), this was also true for models without adjustment for eTIV (eFigure 7B).

### Hippocampal subfields

*Patients with* SZ was associated with lower mean volume in both left and right whole hippocampus as well as in all hippocampal subfields, accompanied with larger right hippocampal fissures. There was increased dispersion in SZ in left whole hippocampus, as well as in left molecular layer, left granule cell layer of the dentate gyrus (GC-DG) and left CA4 (Figure 4A). Models without adjustment for eTIV gave the same results for mean volumes with the exception for left hippocampal fissure, which did not survive correction. Additional dispersion effects were observed for right whole hippocampus, and left CA1 (eFigure 8A). When reanalyzing SZ mean and dispersion models with more stringent exclusion criteria, we obtained similar results (eTable 3). PGR was associated with smaller left CA2/3. None of the subfields showed an association between volume dispersion and PRS (Figure 4B). Models without adjustment for eTIV revealed smaller left and right CA2/3, left granule cell layer of the dentate gyrus (GC-DG), and left CA4 (eFigure 8B) in patients with SZ. Re-analysis with population covariates added to the models, re-analysis with stricter exclusion criteria, and modeling PRS using the first principal component, did not alter conclusions (eTable 4).

## Discussion

In the current study we found that SZ is associated with increased brain heterogeneity in cortical thickness, as well as in cortical volumes, lateral and third ventricles and hippocampal volumes. The findings, based on harmonized analysis protocols for all included datasets, were robust to strict procedures for removing outliers, and follow-up meta-analysis confirmed that multi-site case-control difference cannot be explained by scanning site. These findings are largely in line with a recent meta-analytic study showing increased volumetric heterogeneity in the temporal lobe and lateral and third ventricles^15^. It also extends this meta-analysis by showing increased heterogeneity in cortical thickness as well as in specific hippocampal subfields. Further, increased polygenic risk in healthy individuals was associated with reductions in thickness in frontal and temporal regions, as well as in left dentate gyrus and CA4 and bilateral CA2/3, but not related to thickness dispersion.

We found widespread reductions in cortical thickness in SZ patients, with the characteristic pattern of stronger fronto-temporal effects, as well as global reductions in cortical volume^46^. In addition to these mean changes, we found that SZ is also associated with increased thickness heterogeneity compared to healthy controls. No cortical region showed the opposite pattern of increased homogeneity among patients. As with previous studies we found mean reductions in several brain volumes, with the most robust effects for cortical volume, cerebellum and hippocampus, as well as ventricle enlargement. These regions additionally showed increased heterogeneity in patients compared to controls, and again no region showed increased homogeneity, as might result if a particular region was similarly affected by a common pathophysiological mechanism^15^. Instead, the results are in line with previous studies suggesting substantial neurobiological heterogeneity in SZ^16^. They do however contrast with a recent report^15^ of increased volumetric homogeneity in SZ. One possible explanation is differing sample inclusion criteria, as the previous study included only first-episode psychosis patients. It is not unlikely that an earlier disease stage may offer a more direct window into core aspects of the pathophysiology which later shift towards increased inter-individual variability as patients vary across different illness stages and degrees of severity, as well as differences in treatment and medication status.

PRS reflect cumulative risk across multiple genetic loci, and SZ PRS is associated with several phenotypic traits, including liability for psychiatric disease such as bipolar disorder and schizoaffective disorder, negative symptoms, as well as IQ, working memory performance and brain activation^25,47,48^. SZ-PRS has been associated with cortical gyrification in healthy participants^49^, as well as reductions in global cortical thickness^27^. The current results show that higher SZ-PRS in healthy participants are associated with mean decrease in thickness in fronto-temporal cortices. Interestingly, these shifts in mean thickness were not associated with changes in brain heterogeneity, as was found for patients, pointing to differential effects of genetic risk on mean and heterogeneity changes. A recent study found that SZ environmental risk scores and SZ-PRS scores are both independently related to frontotemporal cortical thinning in patients, but not controls ^27^. This underscores the importance of also investigating environmental risk factors, as well as gene-environment interplay, and their role in explaining the observed clinical and neurobiological heterogeneity

With regard to hippocampal volumes, we found that higher polygenic risk was associated with smaller volumes of the left CA2/3, in the absence of an effect on total hippocampal volume and after correcting for total intracranial volume, suggesting a specific effect of genetic risk on this region. The hippocampus has been hypothesized to play a primary role in the pathophysiology of SZ, through progressive changes to its neural circuits as the disease evolves^29^. Our results also complement recent studies reporting that polygenic risk for SZ is predictive of hippocampal activation during memory encoding^28^, and of polygenic overlap between SZ and hippocampus volume^50^. Also, while the patients showed increased hippocampal heterogeneity, only mean effects were associated with PRS, mirroring the findings on cortical thickness. Thus, the dentate gyrus and CA2/3 emerge as key regions both for the manifestation of, and the genetic risk for, SZ and potentially informative for the classification of sub-types and degrees of severity.

Despite reliable associations between SZ and brain morphometry^14^, PRS is only weakly associated with subcortical volumes, however a recent study found polygenic overlap for SZ and hippocampal, putamen and intracranial volumes^50^. The lack of associations between PRS and subcortical volumes in the current study is in line with most previous reports or PRS^30^.

### Limitations

An important source of heterogeneity in the present case-control sample could be the large number of different scanning-sites included. However, in addition to residualizing for scanner-site in the main analysis, we also performed within-sample analysis and ran meta-analysis, which rule out scanner as a major contributor to the observed effects. Heterogeneity could be associated with differences in medication status and duration of illness. While we did find increased caudate and putamen volumes, indicative of medication effects^51^, these volumes did not show altered heterogeneity in SZ. Still, antipsychotic medication could possibly affect brain heterogeneity in other regions, in absence of a change in mean volume. Investigation of such effects requires carefully controlled settings, and is therefore difficult to address in large-scale multi-site studies. Another possible explanation is that the increased variability is caused by movement artefacts, which are typically greater in clinical populations^52^, however running the analysis in a subset with stricter criteria for dataset exclusion did not alter the conclusions. One important consideration for case-control studies in general is the possibility that healthy controls are abnormally normal compared to the general healthy population due to selection bias and strict exclusion criteria^53^, underscoring the importance of studying the full range of phenotypic variability in the population. Further, the validity of choosing a given p-threshold selection among several possible thresholds when calculating PRS is uncertain. We addressed this by performing follow-up analysis where we performed PCA across PRS calculated across a wide range of thresholds, to derive a more general score of polygenic risk. This approach yielded results converging with the main analysis using a threshold of p < .05. The lack of association between SZ-PRS and brain heterogeneity suggest that the current SZ-PRS scores do not strongly reflect variance-controlling variants. As a composite score, SZ-PRS likely also hides a substantial genetic heterogeneity. A PRS-score calculated using a variance-controlling trait loci (vQTL) approach would likely be more sensitive in detecting such effects. Lastly, SZ is increasingly understood as a neurodevelopmental disorder^54^ and disentangling the sources of heterogeneity in the adult patient population likely requires investigation of the life-span trajectories and aberrant developmental paths.

### Conclusion

There are ongoing efforts to account for brain heterogeneity by means of delineating patient subtypes^8,55^, as well as characterizing patients by their differential degree of affectedness along one or multiple symptom-axes^11,56^. Here we report that SZ is associated with widespread and increased heterogeneity in cortical thickness, and cortical as well as hippocampal volume, beyond the known mean differences, compared controls. The results support to the notion that SZ is a highly heterogeneous disorder, and suggests that important information may be overlooked when only assessing mean differences. In healthy adults SZ-PRS were associated with mean changes in brain areas implicated in SZ, but not associated with altered brain heterogeneity. Taken together, these findings warrant future longitudinal studies which can disentangle the genetic and environmental factors contributing to diverging trajectories and neurobiological heterogeneity, and in particular how these factors contribute to heterogeneity in fronto-temporal cortices and hippocampus.

## Acknowledgements

The authors were funded by the Research Council of Norway (213837, 223273, 229129, 204966/F20, 249795, 251134), the South-Eastern Norway Regional Health Authority (2014097, 2015073, 2016083, 2017112), KG Jebsen Stiftelsen, the European Commission 7th Framework Programme (#602450, IMAGEMEND), the Swedish Research Council (2006-2992, 2006-986, K2007-62X-15077-04-1, 2008-2167, 2008-7573, K2010-62X-15078-07-2, K2012-61X-15078-09-3, 14266-01A,02-03, 2017-949), the regional agreement on medical training and clinical research between Stockholm County Council, the Knut and Alice Wallenbergs Foundation, the HUBIN project, the German Research Foundation DFG (KI 576/14-2, ZI1253/3-1, ZI1253/3-2), Fondazione Con Il Sud and the Hoffmann-La Roche Collaboration. This research has been conducted using the UK Biobank Resource (access code 27412).

This study includes data from several sources. A detailed overview over included cohorts and acknowledgements of their respective funding sources is provided in eTable 1. None of the funding sources played a role in analysis and interpretation of the data; preparation, review, or approval of the manuscript; or decision to submit the manuscript for publication.

## Author contribution statement

D.A. and L.T.W conceived of the study; D.A. performed the analysis with contributions from L.T.W., T.K and D.M; A.C.P., T.M., J.R., F.B., I.A., D.M.B., A.B., C.L.B., S.C., S.D., N.T.D., S.E., H.F.B., L.F., A.D.G., B.H., E.G.J., P.K., M.J.L., A.M.L., G.P., E.S., O.B.S., T.Q., M.Z., and O.A.A contributed with data preprocessing, quality assurance and interpretation of results. D.A., T.K. and L.T.W. wrote the first draft of the paper, and all authors contributed to the final manuscript. D.A. and L.T.W. had full access to all the data in the study and takes responsibility for the integrity of the data and the accuracy of the data analysis.

---

**eFigure 1:**
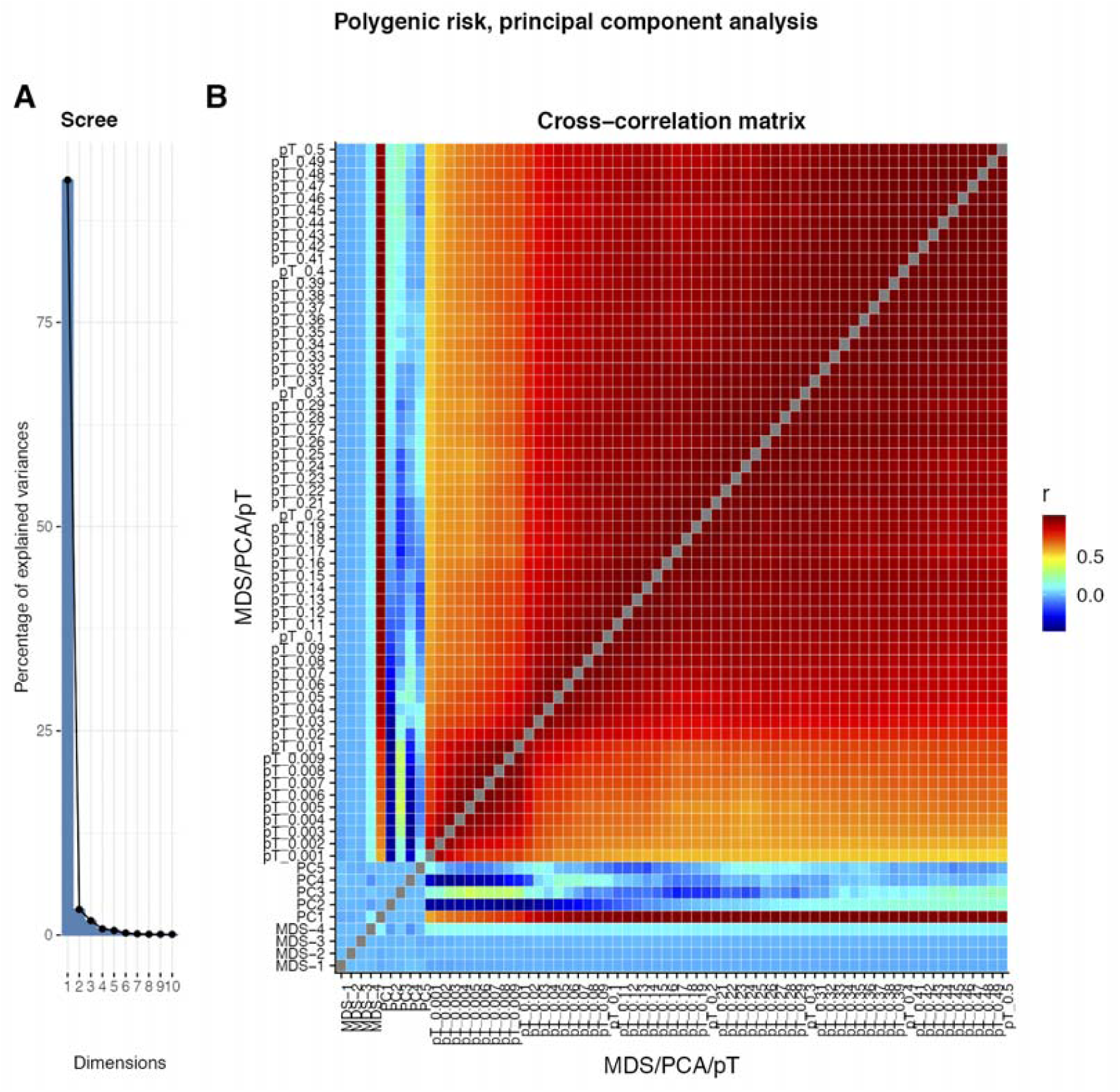
Principal component analysis on PRS scores calculated at different p-thresholds. We performed PCA on PRS scores from the UK Biobank sample (n=12490), calculated from a threshold of p < .001, to p < .5 (500 thresholds, step of .001). **A:** Scree plot showing explained variance for the first 10 components. **B:** Cross-correlation matrix of first four MDS components, first five principal components, and PRS for thresholds from .001 to .5 (steps of .001 up to p < .01, then steps of .01). The first three principal components explained 92.5%, 3.1% and 1.8% of the variance, respectively. The first components showed high correlation with PRS across all thresholds, but relatively lower for the stricter compared to the more lenient p-thresholds. Component 2 correlated negatively with PRS from the stricter p-thresholds, and positive with more lenient p-thresholds. Component 3 was positively correlated with scores calculated with the strictest and most lenient p-thresholds, while correlating negatively with PRS based on the intermediate p-thresholds.

**eFigure 2:**
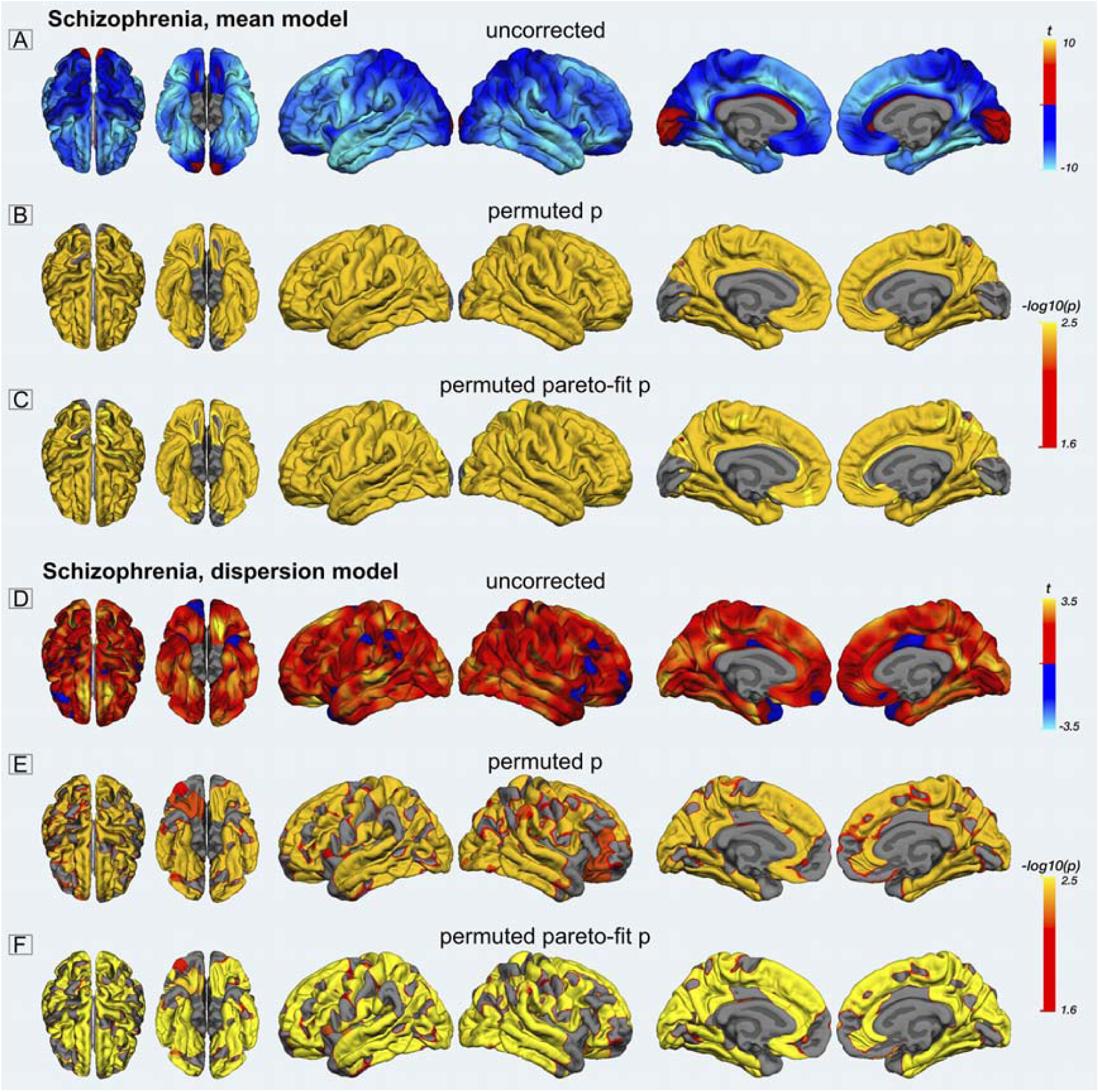
SZ Vertex-wise mean and dispersion models. **A:** unthresholded t-map for the SZ mean model; warm/cold colors represent areas with increased and decreased mean thickness in SZ compared to healthy controls. **B:** -log10(p) map for the SZ-mean model. Thresholded at -log(p) > 1.6 (p < .05, two tailed, FWE). Derived by comparing the vertex-wise values of the actual TFCE-map for the SZ mean-model with TFCE-maps derived from 500 random permutations of the diagnosis labels. **C:** -log10(p) map derived from fitting a generalized Pareto distribution to the tail of the permutation distribution. **D:** unthresholded t-map for the SZ dispersion model; warm/cold colors represent areas with increased and decreased thickness dispersion in SZ compared to healthy controls. **E:** -log10(p) map for the SZ-dispersion model. Thresholded at -log(p) > 1.6 (p < .05, two tailed, FWE). Derived by comparing the vertex-wise values of the actual TFCE-map for the SZ dispersion-model with TFCE-maps derived from 500 random permutations of the diagnosis labels. **F:** -log10(p) map derived from fitting a generalized Pareto distribution to the tail of the permutation distribution.

**eFigure 3:**
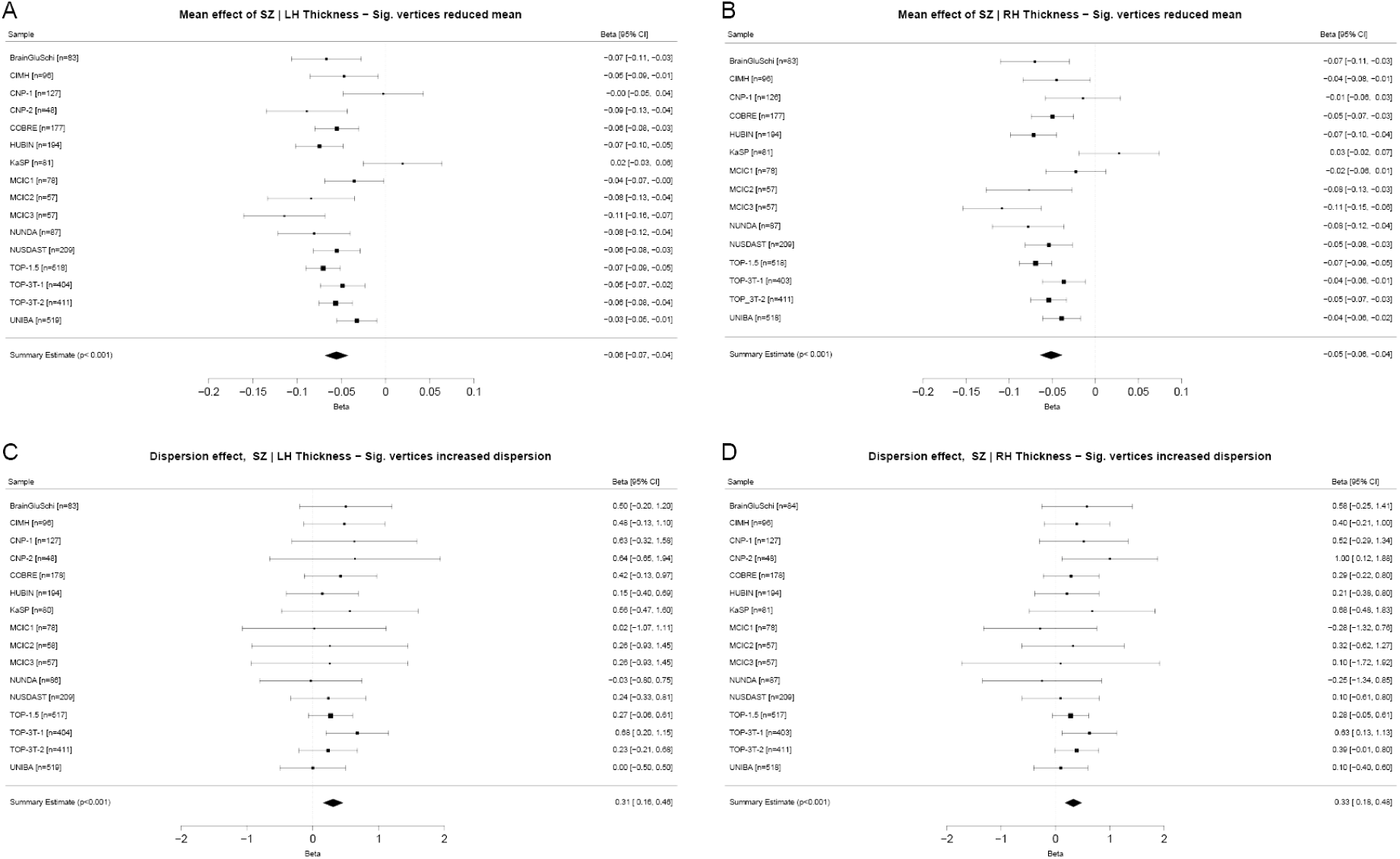
Meta-analysis of SZ mean and dispersion effects with stricter inclusion criteria. The above panel shows the within-sample effects of the mean-model in the vertices showing a significant group-level effect for the left and the right cortical hemisphere. The bottom panel displays the dispersion effects. Both hemispheres showed a significant meta-analytic thickness decrease, as well as a dispersion increase, suggesting that the results from the main analysis are not driven by site-related variance not accounted for in the multi-site regression models. The analysis also included removal of subjects with either eTIV or mean thickness-values at | z | > 3, or significant outlier test (corrected p < .05). For CNP-2, right hemisphere, the DGLM did not converge.

**eFigure 4:**
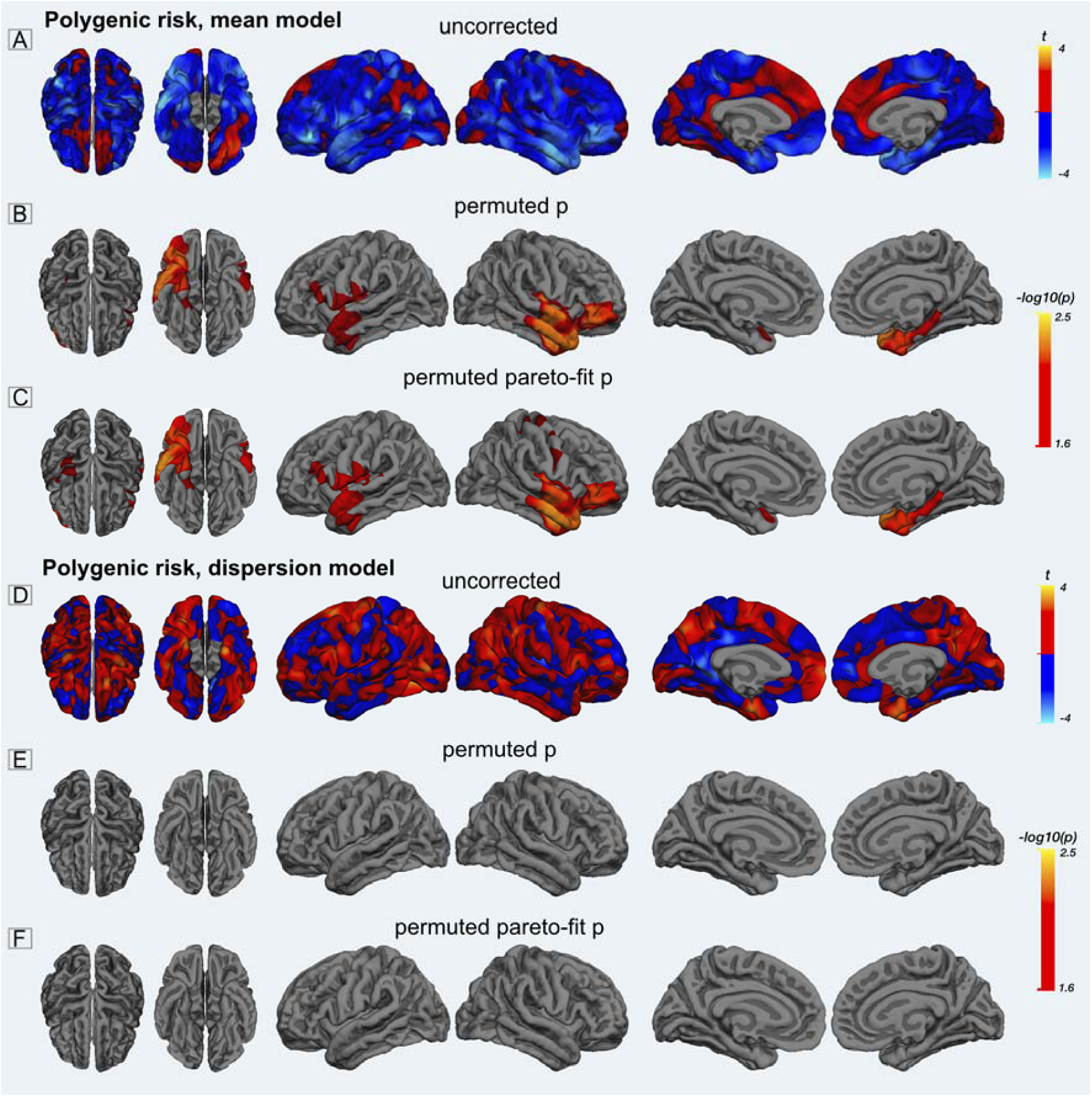
Polygenic risk, vertex-wise mean and dispersion models. **A:** unthresholded t-map for the PRS mean model; warm/cold colors represent areas with increased and decreased mean thickness with increased PRS. **B:** -log10(p) map for the PRS-mean model. Thresholded at -log(p) > 1.6 (p < .05, two tailed, FWE). Derived by comparing the vertex-wise values of the actual TFCE-map for the PRS mean-model with TFCE-maps derived from 500 random permutations of the PRS scores. **C:** -log10(p) map derived from fitting a generalized Pareto distribution to the tail of the permutation distribution. **D:** unthresholded t-map for the PRS dispersion model; warm/cold colors represent areas with increased and decreased thickness dispersion with increased PRS. **E:** -log10(p) map for the PRS-dispersion model. Thresholded at -log(p) > 1.6 (p < .05, two tailed, FWE). Derived by comparing the vertex-wise values of the actual TFCE-map for the SZ dispersion-model with TFCE-maps derived from 500 random permutations of the PRS scores. **F:** -log10(p) map derived from fitting a generalized Pareto distribution to the tail of the permutation distribution.

**eFigure 5:**
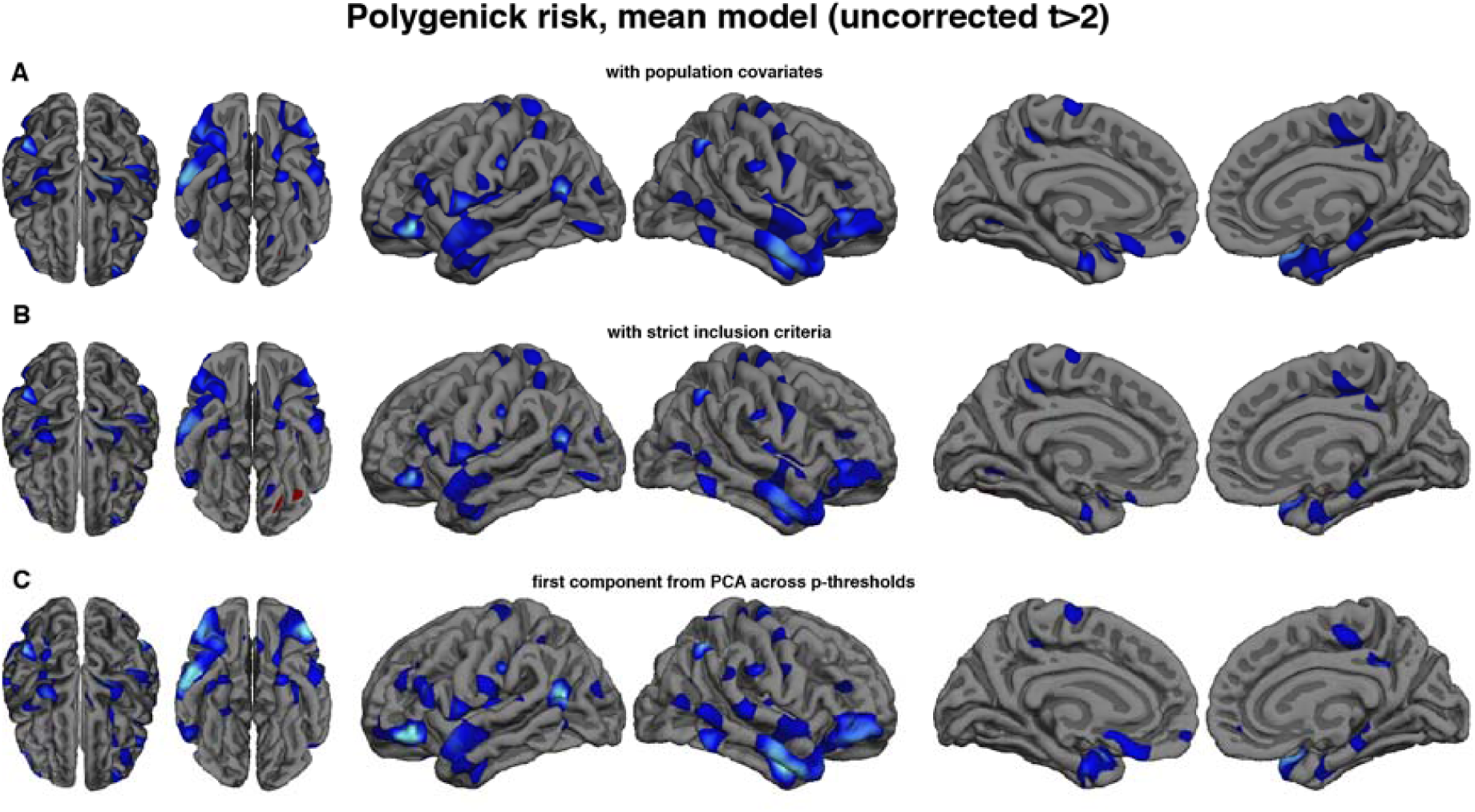
Reanalysis of polygenic risk score models on cortical thickness. To test the robustness of the mean effect of PRS on cortical thickness, we performed several follow-up analysis: **A**: Uncorrected t-map for PRS-scores, including the four first population covariates derived from multidimensional scaling (MDS) analysis. **B:** Uncorrected t-map for reanalysis with extreme scoring individuals removed (mean thickness or eTIV, |z| > 3, or outlier test significant at p < .05, FWE). **C:** Uncorrected t-map for the analysis using the first principal component from the PCA on PRS scores calculated across a wide range of p-thresholds.

**eFigure 6:**
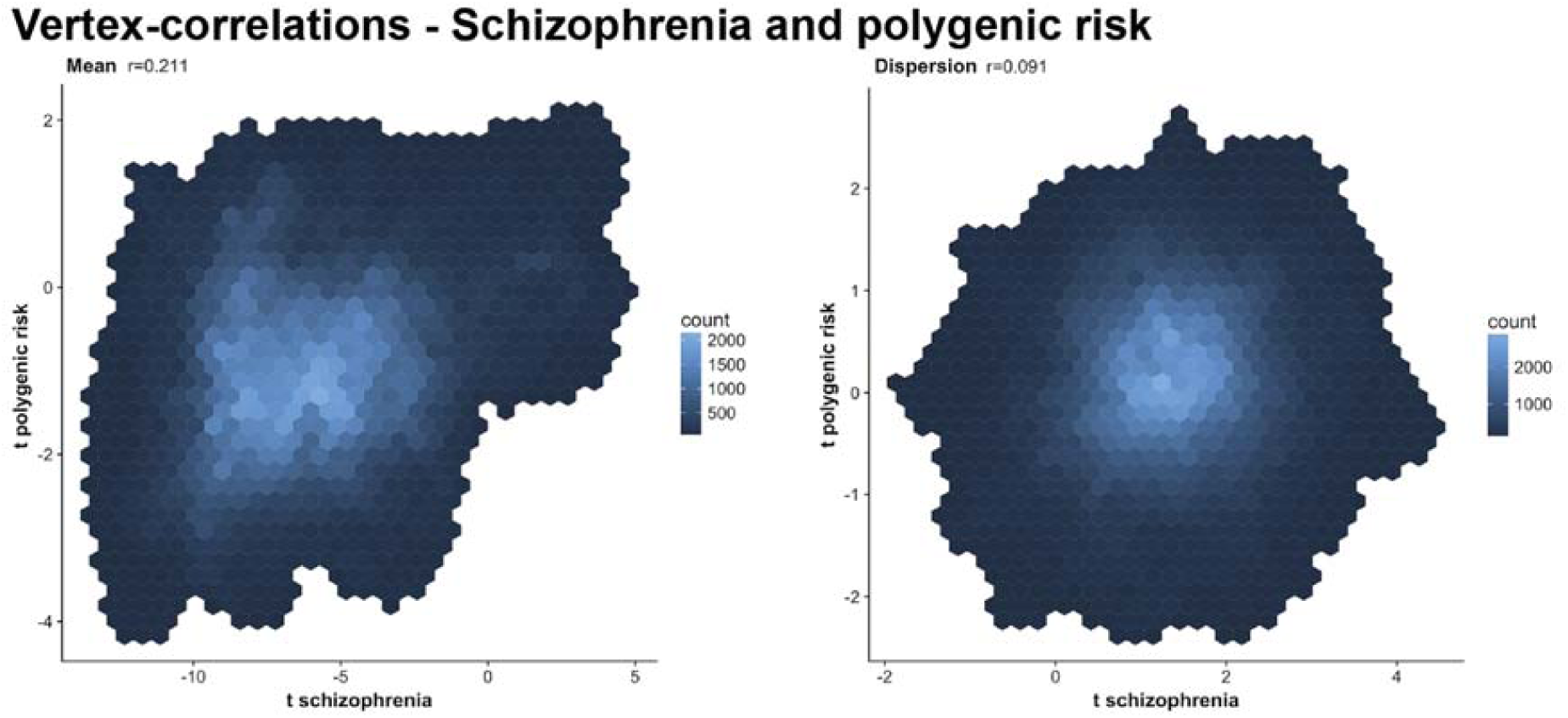
Spatial overlap between SZ and PRS. The vertex-wise t-values for the SZ-models are shown on the x-axis and the vertex-wise t-values for the PRS-models on the Y-axis. The left plot shows the mean-model (vertex-wise correlation of .2) and the right plot shows the dispersion models (vertex-wise correlation of .1). The heatmaps represent the count of vertices at a value.

**eFigure 7:**
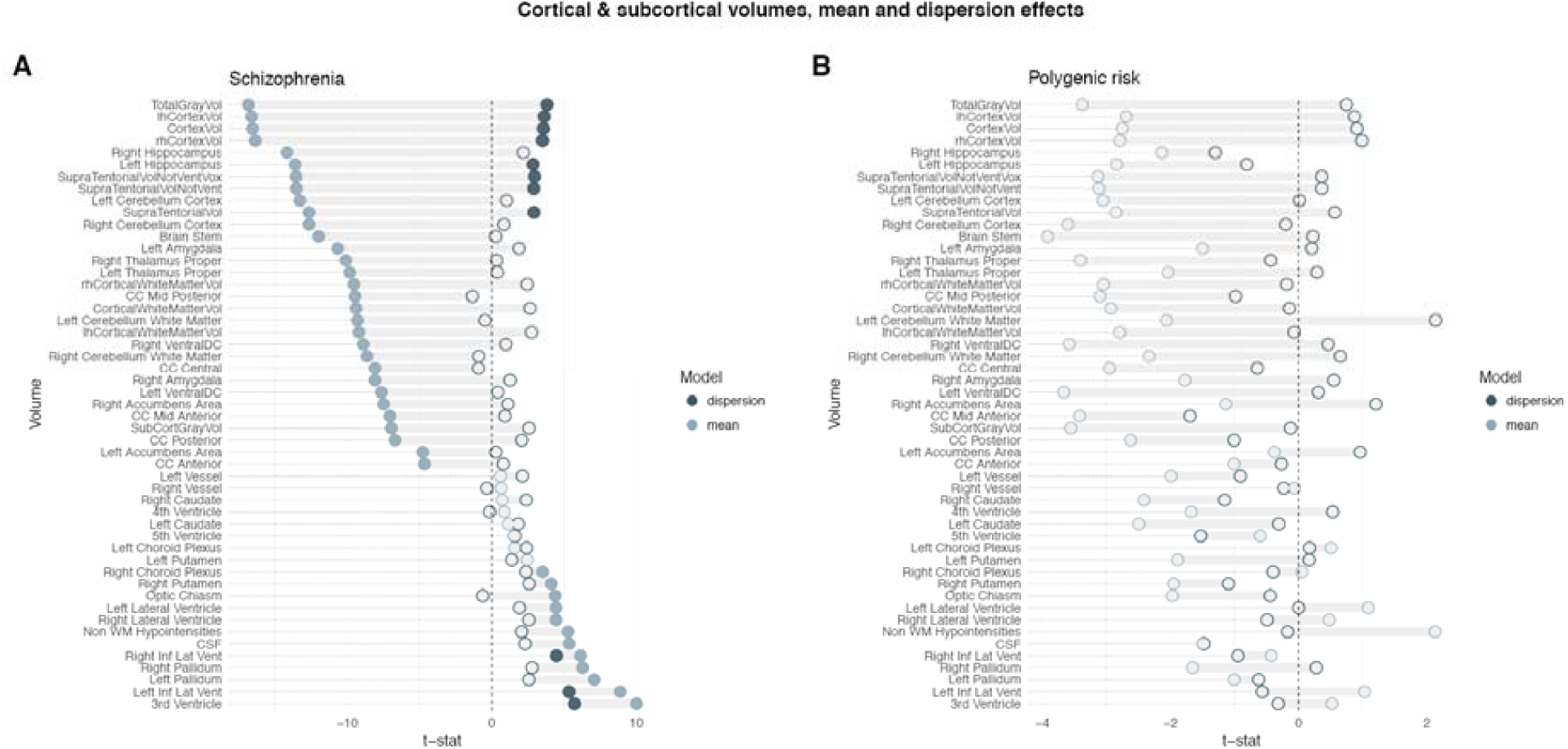
Cortical and subcortical volumes, without adjustment for eTIV. t-statistic for both mean (light blue) and dispersion (dark blue), filled dots mark significant effects after correction for multiple comparisons across regions (5000 permutations, permuted p < .05, FWE). **A:** The SZ-group showed decreased cortical and subcortical volumes, as well as increased ventricles and putamen and pallidum volumes. Cortical, right hippocampal and ventricle volumes were more heterogeneous in the SZ-group compared to healthy controls. **B:** Polygenic risk for SZ was not associated with mean changes nor dispersion in any of the regions.

**eFigure 8:**
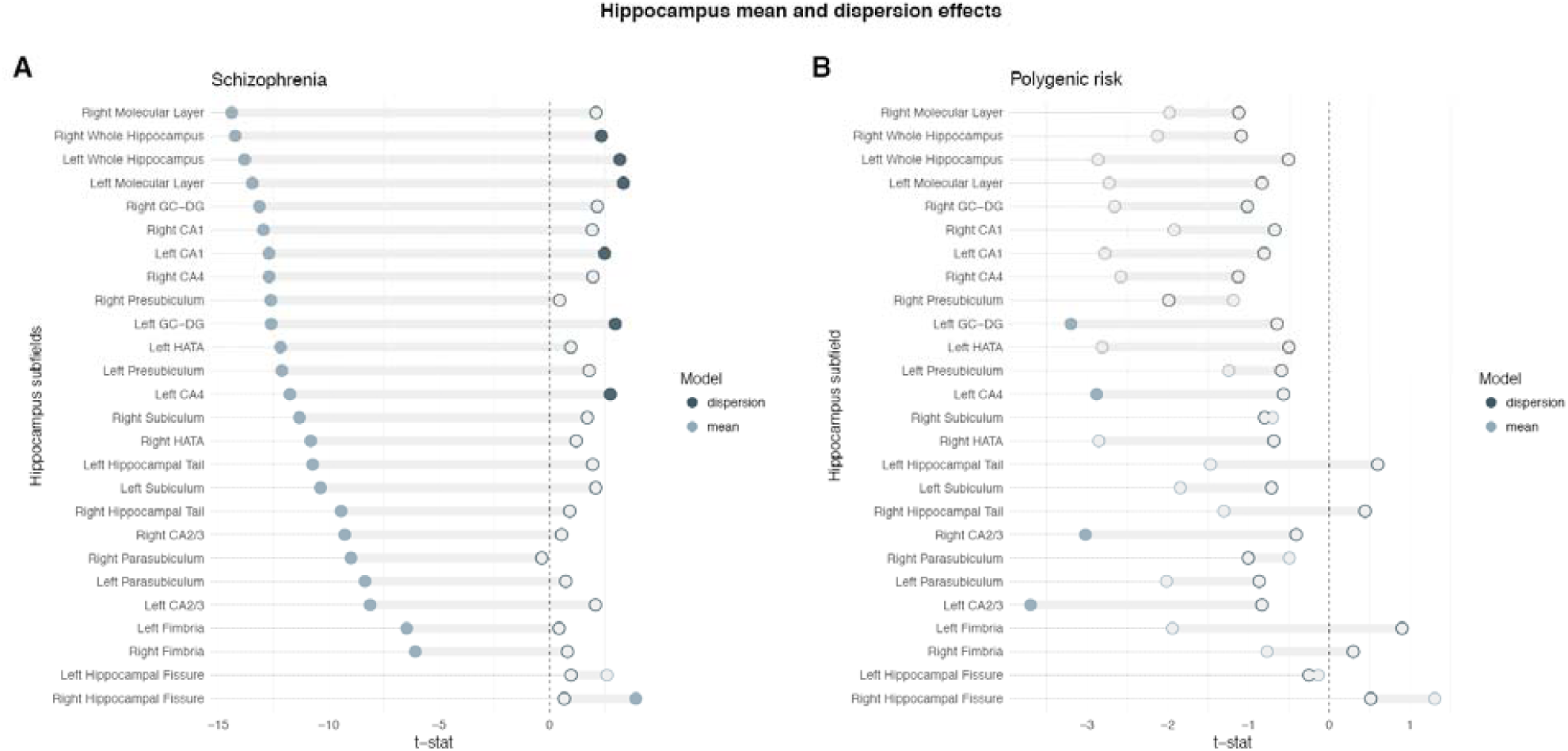
Hippocampus subfields volumes, without adjustment for eTIV. t-statistic for both mean (light blue) and dispersion (dark blue), filled dots mark significant effects after correction for multiple comparisons across regions (5000 permutations, permuted p < .05, FWE). **A:** The SZ-group had decreased hippocampal volumes, and this decrease was also evident in all subfields, accompanied with a decrease of the left hippocampal fissure. Bilateral hippocampal volumes were also more heterogeneous in the SZ-group, and among the subfields this effect was present in the left molecular layer, left CA1, left GC-DG and left CA4. **B:** Polygenic risk for SZ was associated with mean reductions of the left GC-DG, left CA4, and bilateral CA2/3. Neither total hippocampal volumes nor any of the subfields showed a significant association between polygenic risk and volume heterogeneity.

**eTable 1:** Summary of included samples. We included publicly available data as well as data provided by co-authors. Sample reference list contains publications related to the sample as well as data sources.

| Sample | Source | Comment | Reference |
| --- | --- | --- | --- |
| BrainGluSchi | http://schizconnect.org/<br>SchizConnect was funded by NIMH cooperative agreement 1U01 MH097435. | Data obtained from COINS source. Supported by NIMH-grant R01MH084898-01A1. | 1 |
| CIMH | Authors | CIMH was supported by the Deutsche Forschungsgesellschaft (DFG, projects KI 576/14-2, ZI1253/3-1, ZI1253/3-2) and the European Community's Seventh Framework Programme (FP7/2007–2013) grant agreement #602450 (IMAGEMEND) | 2-4 |
| CNP 1 & 2<br>(DS000030) | https://openfmri.org/ | Data sets were obtained from the OpenfMRI database. Data from 2 imaging sites. DS000030 was supported by the Consortium for Neuropsychiatric Phenomics (NIH Roadmap for Medical Research grants UL1-DE019580, RL1MH083268, RL1MH083269, RL1DA024853, RL1MH083270, RL1LM009833, PL1MH083271, and PL1NS062410). | 5,6 |
| COBRE | http://schizconnect.org/ | Data obtained from COINS source. Supported by grant 5P20RR021938 /P20GM103472 from the NIH to Dr. Vince Calhoun. | 7 |
| HUBIN | Authors | HUBIN was supported by the Swedish Research Council (2006-2992, 2006-986, K2007-62X-15077-04-1, 2008-2167, 2008-7573, K2010-62X-15078-07-2, K2012-61X-15078-09-3, 14266-01A,02-03, 2017-949), the regional agreement on medical training and clinical research between Stockholm County Council and the Karolinska Institutet, the Knut and Alice Wallenbergs Foundation and the HUBIN project | 8,9 |
| KaSP | Authors | KaSP was supported by grants from the Swedish Medical Research Council (SE: 2009-7053; 2013-2838; SC: 523-2014-3467), the Swedish Brain Foundation, Ahlén-stiftelsen, Svenska Läkaresamfundet, Petrus och Augusta Hedlunds Stiftelse, Torsten Söderbergs Stiftelse, the AstraZeneca-Karolinska Institutet Joint Research Program in Translational Science, Söderbergs Konungliga Stiftelse, Professor Bror Gadelius Minne, Knut och Alice Wallenbergs stiftelse, Stockholm County Council (ALF and PPG), Centre for Psychiatry Research, KID-funding from the Karolinska Institutet. | 10,11 |
| MCIC-1/2/3 | http://schizconnect.org/ | Data obtained from COINS source. MCIC was supported by the Department of Energy under Award Number DE- FG02-08ER64581. | 12 |
| NUNDA | http://schizconnect.org/ | Data obtained from schizconnect.org. Source: NUNDA Supported by NIMH grant MH056584. | 13 |
| NUSDAST | http://schizconnect.org/ | Data obtained from schizconnect.org. Source: NU_REDCAP. Supported by NIMH Grant 1R01 MH084803 | 14,15 |
| TOP3T 1 & 2, TOP-1.5 | Authors | The work was funded by the Research Council of Norway (213837, 223273, 204966/F20, 213694, 229129, 249795/F20, 248778), the South-Eastern Norway Regional Health Authority (2013-123, 2014-097, 2015-073, #2017-112) and Stiftelsen Kristian Gerhard Jebsen. | 16-19 |
| UKB | https://www.ukbiobank.ac.uk | This research has been conducted using the UK Biobank Resource (access code 27412) | 20 |
| UNIBA | Authors | This work was supported by a “Capitale Umano ad Alta Qualificazione” grant by Fondazione Con Il Sud awarded to Alessandro Bertolino and by a Hoffmann-La Roche Collaboration Grant awarded to Giulio Pergola. This project has received funding from the European Union Seventh Framework Programme for research, technological development and demonstration under grant agreement no. 602450 (IMAGEMEND). This paper reflects only the author's views and the European Union is not liable for any use that may be made of the information contained therein. | 21 |

**eTable 2:**
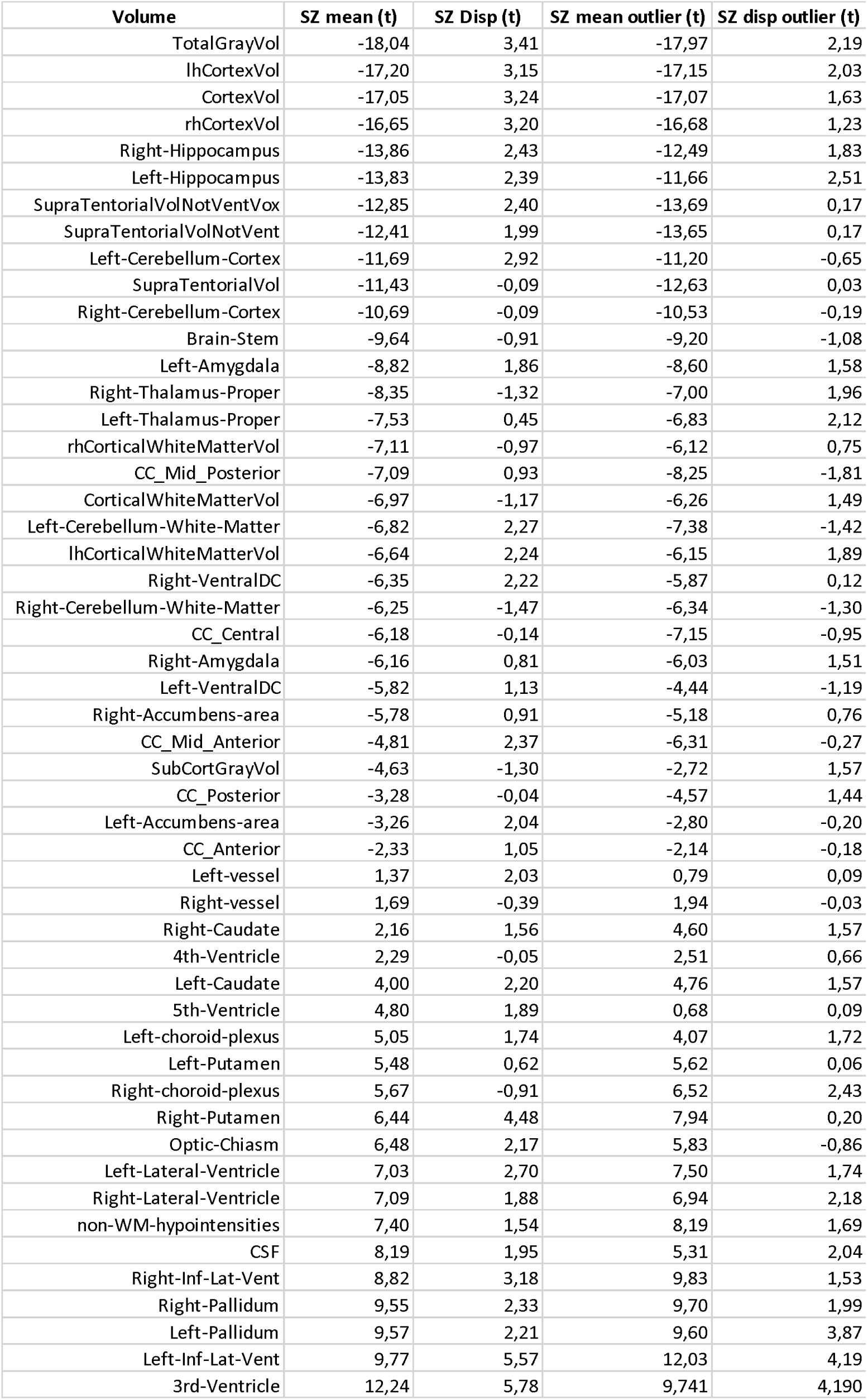
SZ cortical and subcortical analysis with and without outlier removal. We excluded participants based on extreme values (described in Methods) and reran mean and dispersion models without these participants.

**eTable 3:**
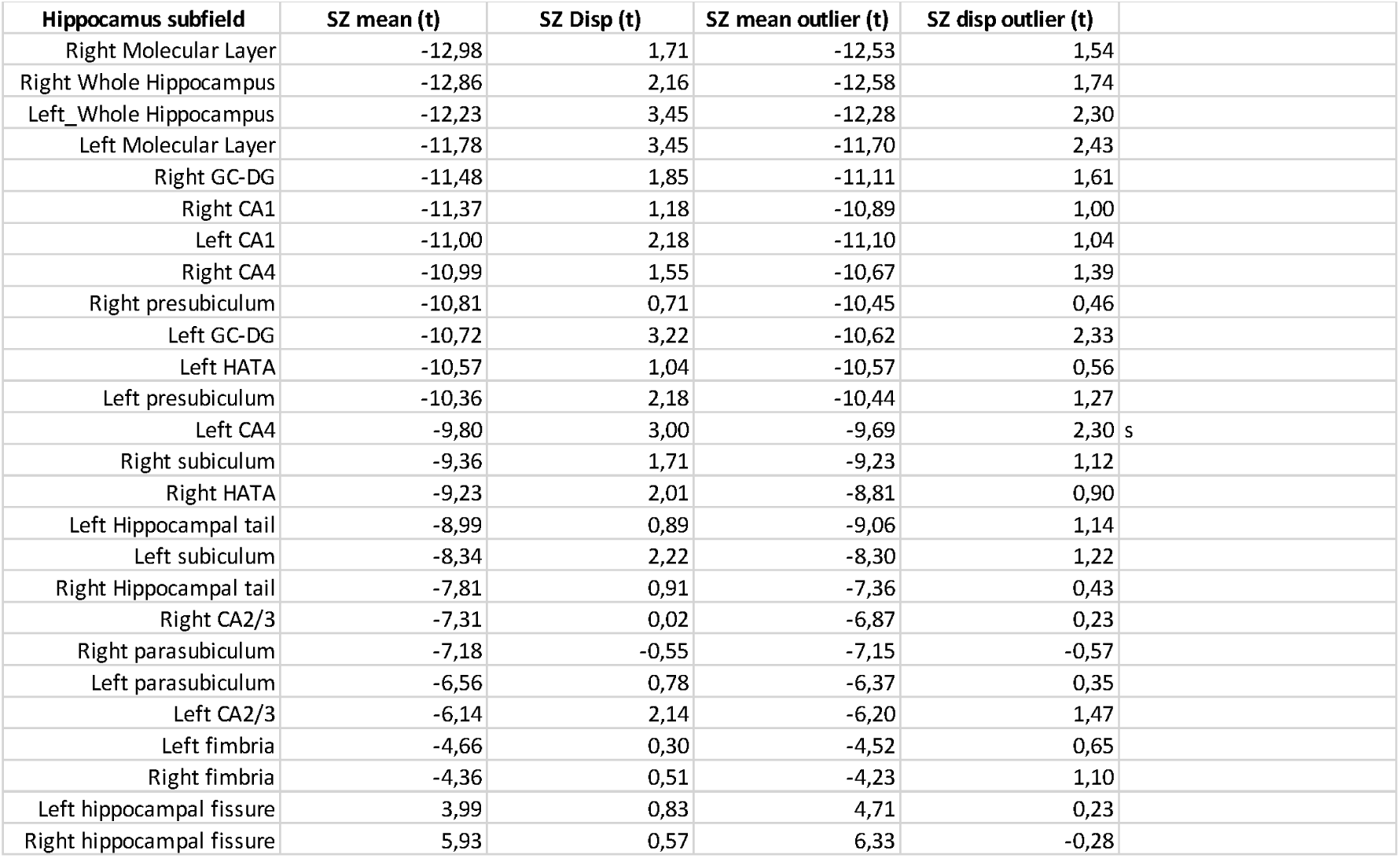
SZ hippocampal subfields analysis with and without outlier removal. We excluded participants based on extreme values (described in Methods) and reran mean and dispersion models without these participants.

**eTable 4:** SZ-PRS hippocampal subfields analysis with and without outlier removal. We excluded participants based on extreme values (described in Methods) and reran mean and dispersion models without these participants.

| Hippocampus subfield | PRS mean (t) | PRS Disp (t) | PRS mean MDS (t) | PRS disp MDS (t) | PRS mean outlier (t) | PRS dis poutlier (t) | PCA1 mean (t) | PCA1 disp (t) |
| --- | --- | --- | --- | --- | --- | --- | --- | --- |
| Left CA2/3 | -3,00 | -1,08 | -2,99 | -0,90 | -2,67 | -1,07 | -2,76 | -0,48 |
| Left GC-DG | -2,26 | -0,93 | -2,15 | -0,70 | -2,10 | -1,43 | -1,95 | -0,12 |
| Right CA2/3 | -2,20 | -0,56 | -2,10 | -0,15 | -1,98 | -0,11 | -2,34 | 0,21 |
| Left CA4 | -1,95 | -0,87 | -1,86 | -0,65 | -1,70 | -1,35 | -1,65 | -0,13 |
| Left Whole hippocampus | -1,83 | -0,61 | -1,62 | -0,34 | -1,68 | -1,12 | -1,36 | -0,19 |
| Right HATA | -2,04 | -0,81 | -1,96 | -0,64 | -2,18 | -0,92 | -2,39 | -0,82 |
| Left HATA | -2,15 | -0,65 | -2,14 | -0,37 | -2,03 | -0,87 | -2,06 | -0,47 |
| Left CA1 | -1,79 | -0,70 | -1,55 | -0,45 | -1,87 | -1,38 | -1,41 | -0,32 |
| Left molecular layer | -1,71 | -0,86 | -1,47 | -0,65 | -1,48 | -1,58 | -1,31 | -0,47 |
| Right GC-DG | -1,61 | -1,18 | -1,39 | -0,86 | -1,59 | -0,87 | -1,51 | -0,43 |
| Right CA4 | -1,54 | -1,31 | -1,33 | -0,94 | -1,44 | -0,78 | -1,45 | -0,63 |
| Right Whole hippocampus | -0,90 | -1,17 | -0,56 | -0,82 | -0,82 | -1,03 | -0,37 | -0,63 |
| Left parasubiculum | -1,37 | -1,13 | -1,35 | -0,88 | -1,09 | -0,70 | -1,50 | -0,65 |
| Right molecular layer | -0,81 | -1,17 | -0,46 | -0,83 | -0,84 | -0,84 | -0,35 | -0,60 |
| Left fimbria | -1,25 | 0,73 | -1,29 | 0,89 | -1,73 | 0,78 | -1,90 | 0,55 |
| Right CA1 | -0,82 | -0,76 | -0,54 | -0,39 | -0,81 | -0,66 | -0,36 | -0,11 |
| Left subiculum | -0,79 | -0,76 | -0,58 | -0,59 | -0,72 | -1,62 | -0,57 | -0,69 |
| Left Hippocampal tail | -0,82 | 0,58 | -0,75 | 0,77 | -0,91 | 0,99 | 0,04 | 0,74 |
| Right Hippocampal tail | -0,53 | 0,47 | -0,41 | 0,70 | -0,46 | 0,48 | 0,66 | 0,42 |
| Left presubiculum | -0,32 | -0,52 | -0,07 | -0,37 | 0,07 | -0,60 | 0,17 | -0,35 |
| Right presubiculum | -0,15 | -1,67 | 0,23 | -1,40 | 0,49 | -1,29 | 0,62 | -0,85 |
| Right fimbria | -0,15 | 0,24 | -0,12 | 0,50 | -0,78 | 0,12 | -0,67 | 0,64 |
| Right subiculum | 0,54 | -0,63 | 0,85 | -0,27 | 0,57 | -0,45 | 0,93 | 0,05 |
| Right parasubiculum | 0,21 | -0,98 | 0,36 | -0,80 | 0,74 | 0,03 | 0,40 | -0,27 |
| Left hippocampal fissure | 0,46 | -0,16 | 0,60 | -0,21 | 0,41 | -0,52 | 0,64 | -0,09 |
| Right hippocampal fissure | 2,12 | 0,73 | 2,30 | 0,83 | 2,19 | 0,88 | 2,35 | 0,60 |

## eMethods

### Samples

For the case-control analysis of cortical thickness and cortical and subcortical volumes we included MRI-scans from 16 cross-sectional study samples with a total of 3161 participants of which 2010 were healthy controls, and 1151 patients diagnosed with SZ. The case-control hippocampus subfield analysis was performed in a sub-sample of 2870 participants. The polygenic risk analysis was performed in a non-overlapping sample consisting of 12490 participants from the UK Biobank (UKB) that had MRI scans and genetic data available, all scanned on the same scanner^35^. All UKB participants diagnosed with any ICD-10 mental or neurological disorder were excluded.

### Genetics

Filtering include removal of SNPs with ambiguous alleles (AT or CG) and of variants in the major histocompatibility complex (MHC; chromosome 6, 26-33Mb), and pruning of SNPs based on linkage disequilibrium (LD) and p-value. All variants in LD with the local SNP with the smallest p-value were removed (the SNP with the smallest p-value within a 250kb window was retained, and all neighbours with a LD r2 > 0.1 were removed; a step known as clumping).

### DGLM

Modeling the dispersion is important for obtaining correct mean parameter estimates if dispersion varies as a function of the predictor, and also allows for systematic investigation into factors associated variability in observations^45^. DGLMs where fitted using the following model specifications for both the mean and dispersion part; case control: Y *∼ Age + Sex + SZ*, and for polygenic risk: Y *∼ Age + Sex + PRS*, where Y is the mean in the first step, and the dispersion in the second step. For the group comparison we then obtain an estimate of the mean difference between groups, as well as an estimate of the difference in dispersion around the mean between SZ and controls for the outcome measure. For the PRS, we obtain an estimate of the linear relationship with PRS and an estimate of the relationship between PRS and the dispersion of the outcome measure.

### Permutation

Due to the computational demand of the vertex-wise analysis (∼5 minutes pr. data chunk, one permutation totals 328 chunks) we ran 600 permutations for each model (case-control, polygenic risk) before fitting a generalized Pareto distribution to the tail of the permutation distribution^47^. For transparency both the raw and fitted permuted p-values are provided.

### Outliers

Participants with a z-score for either eTIV or for volume of interest at |z| > 3 were removed, as well as participants with extreme values based on a statistical outlier-test (Rpackage: car^49^, corrected p < .05 for the linear models *Y ∼ Age + Sex + SZ* and *Y ∼ Age + Sex + PRS)*.

### PRS-PCA

PCA was performed on SZ-PRS scores from p-thresholds p < .001 to p < −05 (500 thresholds, steps of .001), before extracting subject loadings from the first three principal components for associations with cortical thickness. The first three principal components explained 92.5%, 3.1% and 1.8% of the variance, respectively (eFigure 1A). The first component (PRS-PC1) showed high correlation with PRS across all thresholds, but relatively lower correlations for the PRS calculated at the stricter p-thresholds compared to the PRS calculated using more lenient p-thresholds. Component 2 (PRS-PC2) correlated negatively with PRS from the stricter p-thresholds, and positive with more lenient p-thresholds. Component 3 (PRS-PC3) was positively correlated with scores calculated with the strictest and most lenient p-thresholds, while correlating negatively with PRS based on the intermediate p-thresholds (eFigure 1B).

## References

1. WHO. The Global Burden of Disease: 2004 Update. In: WHO Press (2008).

2. Lakhan SE, Vieira KF. Schizophrenia pathophysiology: are we any closer to a complete model? Annals of General Psychiatry. 2009;8(1):12.

3. Owen MJ, Sawa A, Mortensen PB. Schizophrenia. The Lancet. 2016;388(10039):86–97.

4. Van Rheenen TE, Lewandowski KE, Tan EJ, et al. Characterizing cognitive heterogeneity on the schizophrenia–bipolar disorder spectrum. Psychological Medicine. 2017;47(10):1848– 1864.

5. Malhotra AK. Dissecting the Heterogeneity of Treatment Response in First-Episode Schizophrenia. Schizophrenia Bulletin. 2015;41(6):1224–1226.

6. Huber G. The heterogeneous course of schizophrenia. Schizophrenia Research. 1997;28(2):177–185.

7. Weinberg D, Lenroot R, Jacomb I, et al. Cognitive subtypes of schizophrenia characterized by differential brain volumetric reductions and cognitive decline. JAMA Psychiatry. 2016;73(12):1251–1259.

8. Zhang T, Koutsouleris N, Meisenzahl E, Davatzikos C. Heterogeneity of Structural Brain Changes in Subtypes of Schizophrenia Revealed Using Magnetic Resonance Imaging Pattern Analysis. Schizophrenia Bulletin. 2015;41(1):74–84.

9. Sugihara G, Oishi N, Son S, Kubota M, Takahashi H, Murai T. Distinct Patterns of Cerebral Cortical Thinning in Schizophrenia: A Neuroimaging Data-Driven Approach. Schizophrenia Bulletin. 2017;43(4):900–906.

10. Seaton BE, Goldstein G, Allen DN. Sources of Heterogeneity in Schizophrenia: The Role of Neuropsychological Functioning. Neuropsychology review. 2001;11(1):45–67.

11. Koutsouleris N, Gaser C, Jäger M, et al. Structural correlates of psychopathological symptom dimensions in schizophrenia: A voxel-based morphometric study. NeuroImage. 2008;39(4):1600–1612.

12. van Erp TGM, Walton E, Hibar DP, et al. Cortical brain abnormalities in 4474 individuals with schizophrenia and 5098 controls via the ENIGMA consortium. Biological Psychiatry. 2018.

13. Moberget T, Doan NT, Alnæs D, et al. Cerebellar volume and cerebellocerebral structural covariance in schizophrenia: a multisite mega-analysis of 983 patients and 1349 healthy controls. Molecular psychiatry. 2017.

14. van Erp TGM, Hibar DP, Rasmussen JM, et al. Subcortical brain volume abnormalities in 2028 individuals with schizophrenia and 2540 healthy controls via the ENIGMA consortium. Molecular psychiatry. 2015;21:547.

15. Brugger SP, Howes OD. Heterogeneity and homogeneity of regional brain structure in schizophrenia: A meta-analysis. JAMA Psychiatry. 2017;74(11):1104–1111.

16. Wolfers T, Doan NT, Kaufmann T, et al. Extensive interindividual differences in schizophrenia and bipolar disorder: mapping biological heterogeneity in reference to normative brain ageing. JAMA psychiatry. in press.

17. Wolfers T, Buitelaar JK, Beckmann CF, Franke B, Marquand AF. From estimating activation locality to predicting disorder: A review of pattern recognition for neuroimaging-based psychiatric diagnostics. Neuroscience and biobehavioral reviews. 2015;57:328–349.

18. Doan NT, Kaufmann T, Bettella F, et al. Distinct multivariate brain morphological patterns and their added predictive value with cognitive and polygenic risk scores in mental disorders. NeuroImage: Clinical. 2017;15:719–731.

19. Gopal S, Miller RL, Michael A, et al. Spatial Variance in Resting fMRI Networks of Schizophrenia Patients: An Independent Vector Analysis. Schizophrenia Bulletin. 2016;42(1):152–160.

20. Gopal S, Miller RL, Baum SA, Calhoun VD. Approaches to Capture Variance Differences in Rest fMRI Networks in the Spatial Geometric Features: Application to Schizophrenia. Frontiers in neuroscience. 2016;10:85.

21. Sullivan PF, Kendler KS, Neale MC. Schizophrenia as a complex trait: Evidence from a meta-analysis of twin studies. Archives of General Psychiatry. 2003;60(12):1187–1192.

22. Schizophrenia Working Group of the Psychiatric Genomics C. Biological insights from 108 schizophrenia-associated genetic loci. Nature. 2014;511(7510):421–427.

23. The International Schizophrenia C. Common polygenic variation contributes to risk of schizophrenia and bipolar disorder. Nature. 2009;460:748.

24. Jones HJ, Stergiakouli E, Tansey KE, et al. Phenotypic manifestation of genetic risk for schizophrenia during adolescence in the general population. JAMA Psychiatry. 2016;73(3):221–228.

25. Kauppi K, Westlye LT, Tesli M, et al. Polygenic Risk for Schizophrenia Associated With Working Memory-related Prefrontal Brain Activation in Patients With Schizophrenia and Healthy Controls. Schizophrenia Bulletin. 2015;41(3):736–743.

26. Walton E, Geisler D, Lee PH, et al. Prefrontal Inefficiency Is Associated With Polygenic Risk for Schizophrenia. Schizophrenia Bulletin. 2014;40(6):1263–1271.

27. Neilson E, Bois C, Gibson J, et al. Effects of environmental risks and polygenic loading for schizophrenia on cortical thickness. Schizophrenia Research. 2017;184:128–136.

28. Chen Q, Ursini G, Romer AL, et al. Schizophrenia polygenic risk score predicts mnemonic hippocampal activity. Brain : a journal of neurology. 2018;141(4):1218–1228.

29. Lieberman JA, Girgis RR, Brucato G, et al. Hippocampal dysfunction in the pathophysiology of schizophrenia: a selective review and hypothesis for early detection and intervention. Molecular psychiatry. 2018.

30. Reddaway JT, Doherty JL, Lancaster T, Linden D, Walters JT, Hall J. Genomic and Imaging Biomarkers in Schizophrenia. In: Current Topics in Behavioral Neurosciences. Berlin, Heidelberg: Springer Berlin Heidelberg; 2018:1–28.

31. Fraser HB, Schadt EE. The Quantitative Genetics of Phenotypic Robustness. PloS one. 2010;5(1):e8635.

32. Dwyer DB, Cabral C, Kambeitz-Ilankovic L, et al. Brain Subtyping Enhances The Neuroanatomical Discrimination of Schizophrenia. Schizophrenia Bulletin. 2018:sby008– sby008.

33. Conley D, Johnson R, Domingue B, Dawes C, Boardman J, Siegal M. A sibling method for identifying vQTLs. PloS one. 2018;13(4):e0194541.

34. Fischl B. FreeSurfer. Neuroimage. 2012;62(2):774–781.

35. Fischl B, Salat DH, Busa E, et al. Whole brain segmentation: automated labeling of neuroanatomical structures in the human brain. Neuron. 2002;33(3):341–355.

36. Dale AM, Fischl B, Sereno MI. Cortical surface-based analysis. I. Segmentation and surface reconstruction. Neuroimage. 1999;9(2):179–194.

37. Fischl B, Sereno MI, Dale AM. Cortical surface-based analysis. II: Inflation, flattening, and a surface-based coordinate system. Neuroimage. 1999;9(2):195–207.

38. Iglesias JE, Augustinack JC, Nguyen K, et al. A computational atlas of the hippocampal formation using ex vivo, ultra-high resolution MRI: Application to adaptive segmentation of in vivo MRI. (1095-9572 (Electronic)).

39. Euesden J, Lewis CM, O’Reilly PF. PRSice: Polygenic Risk Score software. Bioinformatics. 2015;31(9):1466–1468.

40. Schizophrenia Working Group of the Psychiatric Genomics C. Biological insights from 108 schizophrenia-associated genetic loci. Nature. 2014;511:421.

41. Wood S. mgcv: Mixed GAM Computation Vehicle with Automatic Smoothness Estimation (v 1.8-23). https://CRAN.R-project.org/package=mgcv. 2018.

42. Dunn PK, Smyth GK. dglm: Double Generalized Linear Models (v. 1.8.3). https://CRAN.R-project.org/package=dglm. 2016.

43. Winkler AM, Ridgway GR, Webster MA, Smith SM, Nichols TE. Permutation inference for the general linear model. NeuroImage. 2014;92:381–397.

44. Viechtbauer W. Conducting meta-analyses in R with the metafor. Journal of Statistical Software. 2010;36(3): http://www.jstatsoft.org/v36/i03/.

45. Kassambara A, Mundt F. factoextra: Extract and Visualize the Results of Multivariate Data Analyses. R package (v. 1.0.5) https://CRAN.R-project.org/package=factoextra. 2017.

46. van Erp TGM, Walton E, Hibar DP, et al. Cortical Brain Abnormalities in 4474 Individuals With Schizophrenia and 5098 Control Subjects via the Enhancing Neuro Imaging Genetics Through Meta Analysis (ENIGMA) Consortium. Biological Psychiatry. 2018.

47. Mistry S, Harrison JR, Smith DJ, Escott-Price V, Zammit S. The use of polygenic risk scores to identify phenotypes associated with genetic risk of schizophrenia: Systematic review. Schizophrenia Research. 2018;197:2–8.

48. Tesli M, Espeseth T, Bettella F, et al. Polygenic risk score and the psychosis continuum model. Acta Psychiatrica Scandinavica. 2014;130(4):311–317.

49. Liu B, Zhang X, Cui Y, et al. Polygenic Risk for Schizophrenia Influences Cortical Gyrification in 2 Independent General Populations. Schizophrenia Bulletin. 2017;43(3):673– 680.

50. Smeland OB, Wang Y, Frei O, et al. Genetic Overlap Between Schizophrenia and Volumes of Hippocampus, Putamen, and Intracranial Volume Indicates Shared Molecular Genetic Mechanisms. Schizophrenia Bulletin. 2018;44(4):854–864.

51. Jørgensen KN, Nesvåg R, Gunleiksrud S, Raballo A, Jönsson EG, Agartz I. First-and second-generation antipsychotic drug treatment and subcortical brain morphology in schizophrenia. European Archives of Psychiatry and Clinical Neuroscience. 2016;266(5):451–460.

52. Reuter M, Tisdall MD, Qureshi A, Buckner RL, van der Kouwe AJW, Fischl B. Head motion during MRI acquisition reduces gray matter volume and thickness estimates. NeuroImage. 2015;107:107–115.

53. Schwartz S, Susser E. The use of well controls: an unhealthy practice in psychiatric research. Psychological Medicine. 2011;41(6):1127–1131.

54. Insel TR. Rethinking schizophrenia. Nature. 2010;468(7321):187–193.

55. Honnorat N, Dong A, Meisenzahl-Lechner E, Koutsouleris N, Davatzikos C. Neuroanatomical heterogeneity of schizophrenia revealed by semi-supervised machine learning methods. Schizophrenia Research. 2017.

56. Viher PV, Stegmayer K, Giezendanner S, et al. Cerebral white matter structure is associated with DSM-5 schizophrenia symptom dimensions. NeuroImage: Clinical. 2016;12:93–99.

